# AgRP neuron activity promotes associations between sensory and nutritive signals to guide flavor preference

**DOI:** 10.1101/2023.09.19.558483

**Authors:** Nathaniel T. Nyema, Aaron D. McKnight, Alexandra G. Vargas-Elvira, Heather M. Schneps, Elizabeth G. Gold, Kevin P. Myers, Amber L. Alhadeff

## Abstract

**Objective:** The learned associations between sensory cues (e.g., taste, smell) and nutritive value (e.g., calories, post-ingestive signaling) of foods powerfully influences our eating behavior [1], but the neural circuits that mediate these associations are not well understood. Here, we examined the role of agouti-related protein (AgRP)-expressing neurons – neurons which are critical drivers of feeding behavior [2; 3] – in mediating flavor-nutrient learning (FNL).

**Methods:** Because mice prefer flavors associated with AgRP neuron activity suppression [4], we examined how optogenetic stimulation of AgRP neurons during intake influences FNL, and used fiber photometry to determine how endogenous AgRP neuron activity tracks associations between flavors and nutrients.

**Results:** We unexpectedly found that tonic activity in AgRP neurons during FNL potentiated, rather than prevented, the development of flavor preferences. There were notable sex differences in the mechanisms for this potentiation. Specifically, in male mice, AgRP neuron activity increased flavor consumption during FNL training, thereby strengthening the association between flavors and nutrients. In female mice, AgRP neuron activity enhanced flavor-nutrient preferences independently of consumption during training, suggesting that AgRP neuron activity enhances the reward value of the nutrient-paired flavor. Finally, *in vivo* neural activity analyses demonstrated that acute AgRP neuron dynamics track the association between flavors and nutrients in both sexes.

**Conclusions:** Overall, these data (1) demonstrate that AgRP neuron activity enhances associations between flavors and nutrients in a sex-dependent manner and (2) reveal that AgRP neurons track and update these associations on fast timescales. Taken together, our findings provide new insight into the role of AgRP neurons in assimilating sensory and nutritive signals for food reinforcement.

## 1. Introduction

For most people, the pleasure that we get from food contributes to our eating behaviors. What makes food rewarding? It is increasingly appreciated that both sensory (e.g., taste, smell) [5] and nutritive (e.g., caloric content) [6–8] properties contribute to the reinforcing value of food. These properties do not act independently; in fact, learned associations between these sensory and nutritive food components robustly drive food preferences [1]. This is especially relevant in our modern food environment, where we are constantly exposed to sensory cues that predict tasty, energy-dense foods. Because of global increases in the prevalence of obesity and related metabolic diseases [9], understanding the neural basis of the association between sensory and nutritive food properties, and how it contributes to food preference, is especially urgent and important.

The process of associating sensory cues with nutrient content is called flavor-nutrient learning (FNL) and can be studied in the laboratory by pairing arbitrary flavors with post-ingestive nutrients. This paradigm has been applied to rodent models to gain insight into the development of food preferences [1]. These studies have revealed that rodents learn to prefer flavors associated with gut detection of nutritive over nonnutritive foods.

Signaling by sensory and nutritive cues originates in the periphery and converges in the brain. What are the central circuits that mediate FNL? There is both pharmacological and physiological evidence that FNL involves dopamine signaling in several brain regions [10] and is dependent on vagal signaling from the gut to the hindbrain [11; 12], although there is also evidence for vagal-independent pathways [13–15]. But aside from the role of this gut-brain reward circuitry [7; 16], the contributions of other feeding circuits to FNL remain largely unexplored.

One potential contributor to FNL is the population of hypothalamic agouti-related protein (AgRP)-expressing neurons. Activity in these neurons is both necessary and sufficient to drive feeding behavior [2; 3; 17]. Further, *in vivo* neural activity recordings have demonstrated that these neurons are highly active during food deprivation and inhibited by both sensory food cues (transient inhibition) [4; 18; 19] and post-ingestive nutrient signaling (sustained inhibition) [20; 21]. Interestingly, mice prefer flavors that are associated with the suppression of AgRP neuron activity and avoid flavors that are associated with elevated AgRP neuron activity during consumption [4]. However, elevated AgRP neuron activity *prior to consumption* increases the preference for an associated flavor [22]. Further, stimulation of AgRP neurons potentiates the dopamine response to caloric food [23]. These data suggest that while AgRP neuron activity transmits negative valence in the absence of food, the suppression of AgRP activity that results from nutrient ingestion may contribute to the reinforcing value of nutrients. Together, these findings led us to hypothesize that AgRP neuron activity contributes to the development of FNL.

Here, we performed a series of experiments to determine whether and how activity in AgRP neurons contributes to, and/or is modulated by, FNL. Because mice prefer flavors associated with AgRP neuron activity reduction [4], we (1) tested whether inhibition of AgRP neuron activity that would ordinarily result from gut nutrient sensing is necessary for FNL, and (2) examined endogenous AgRP neuron activity dynamics during the development (training) and expression (testing) of flavor preferences. Overall, our results unexpectedly demonstrate that AgRP neuron activity strengthens rather than prevents FNL via different mechanisms in male and female mice, and reveal their role in tracking short-timescale associations between flavors and nutrients.

## 2. Results

### 2.1 AgRP neuron stimulation potentiates flavor-nutrient learning

Mice prefer flavors associated with reduced AgRP neuron activity [4], suggesting that the suppression of AgRP neuron activity by food may be critical for the development of food preferences. We therefore tested whether AgRP neuron activity suppression is necessary for the development of a preference for nutrient-associated flavors. To do so, we optogenetically stimulated AgRP neurons in hungry mice during FNL training sessions to block reductions in AgRP neuron activity by post-ingestive nutrients (Figure 1A). We engineered *Agrp-Ires-Cre* mice to express channelrhodopsin-2 (ChR2, by crossing with *Ai32* mice) or tdTomato (control, by crossing with *Ai9* mice) to enable optogenetic stimulation of AgRP neurons, and implanted them with chronic gastric catheters for direct delivery of nutrients to the stomach. As previously shown [2; 3], ChR2-mediated activation of AgRP neurons in *ad libitum*-fed mice increased Fos expression (Figure 1B) and stimulated food intake (Figure 1C, t(13) = −13.857, p<0.0001). We food restricted the mice and trained them in an FNL protocol adapted from previous studies (Figure 1D, S1A) [24–26]. Briefly, mice were given access to two flavors in alternating sessions; licking for the flavor designated as the conditioned stimulus (CS)+ triggered a contemporaneous gastric infusion of glucose, whereas licking the CS- flavor triggered a water infusion (Figure 1D). Mice receiving AgRP neuron stimulation consumed significantly more of both flavors during training (Figure S1B, S1C, Chr2: F(1, 27)=14.058, p<0.001). Further, all mice (regardless of stimulation condition) licked more for the CS+ compared to CS- flavor during training (Figure S1B, S1C, CS: F(1,27)=45.376, p<0.0001), which is consistent with prior work and verifies that mice were responsive to gut nutrient sensing. Contrary to our hypothesis, we found that preventing AgRP neuron inhibition did not block FNL (Figure 1E-1G). In fact, 2-bottle choice tests between the CS+ and CS- flavors in the absence of gastric infusion revealed that while AgRP neuron stimulation in hungry mice during FNL training did not increase overall CS+ and CS- licks during testing (Figure 1F, t(27) = 1.853, p=ns) it significantly *increased* preference for the CS+ over CS- flavor (Figure 1G, U=152, p<0.05). Therefore, stimulating AgRP neurons in hungry mice during FNL training, thereby preventing nutrient-mediated suppression of AgRP neurons, enhances preferences for nutrient-associated flavors.

**Figure 1.**
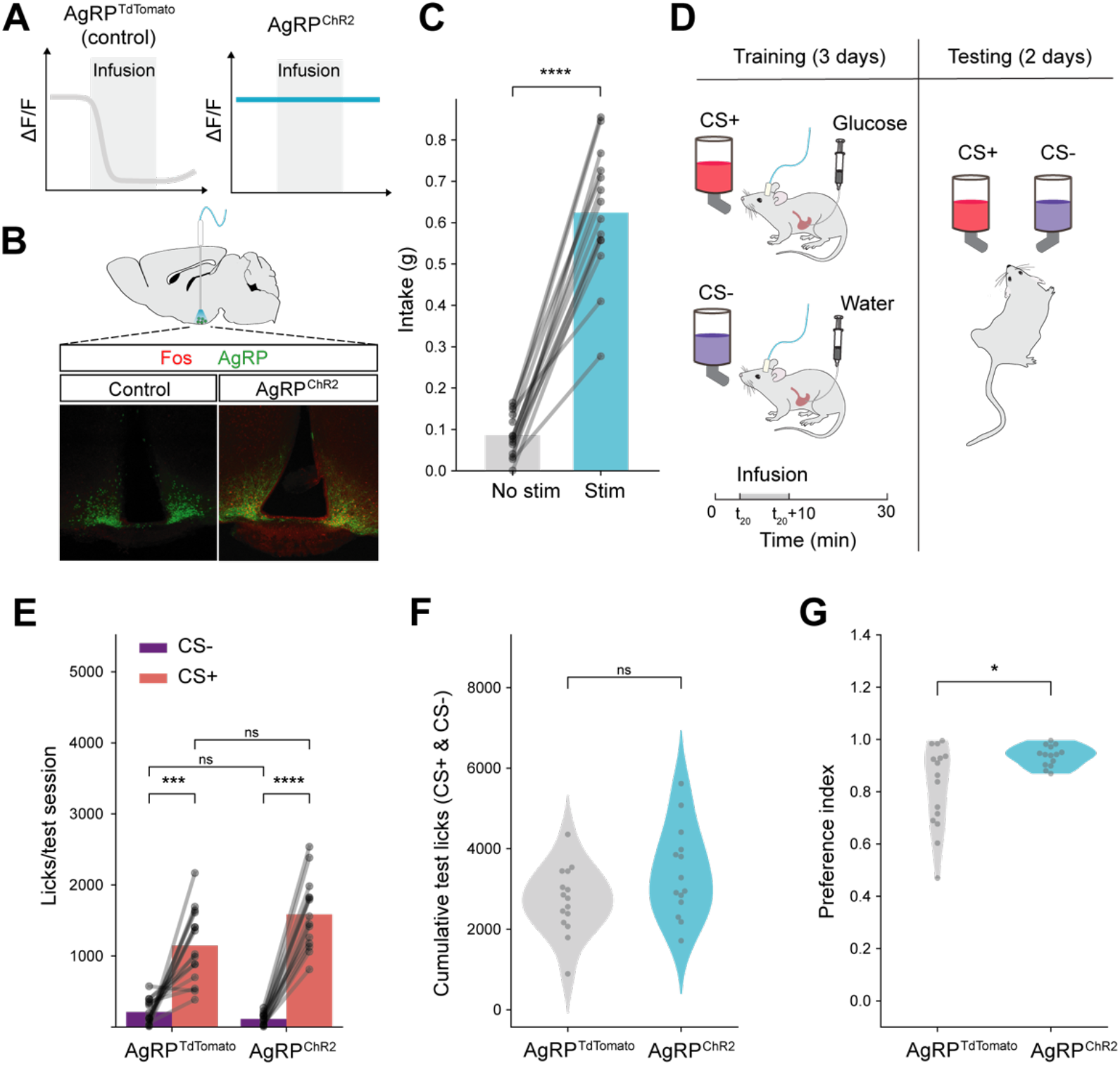
AgRP neuron activity potentiates flavor-nutrient learning. (A) Schematic for AgRP neuron stimulation: optogenetic activation of AgRP neurons (AgRP^ChR2^) was used to prevent inhibition of AgRP neurons during infusion of nutrients (glucose) in (D). (B) Schematic of brain stimulation and representative images of Fos expression in control and AgRP^ChR2^ mice. (C) Food intake in AgRP^ChR2^ mice with and without optogenetic stimulation (n=14). Paired t-test, t(13) = −13.857, p<0.0001. Bar graph represents mean. (D) Simplified schematic of flavor-nutrient learning (FNL) protocol. Mice received a 10-min gastric infusion of nutrients (glucose) or water triggered by 20 flavor licks (at time=t_20_). See Figure S1 and Methods for detailed protocol. (E) Average number of licks per test session following FNL protocol in control (AgRP^tdTomato^) (n=15) and experimental (AgRP^ChR2^) (n=14) mice. Mixed ANOVA, Chr2: F(1,27)=3.506, p=ns; CS: F(1,27)=132.518, p<0.0001; Chr2*CS: F(1,27)=6.673, p<0.05. (F) Total number of CS+ and CS- licks in test sessions in control (AgRP^tdTomato^) (n=15) and experimental (AgRP^ChR2^) (n=14) mice. T-test, t(27) = 1.853, p=ns. (G) Preference index (proportion of CS+ licks during testing, see Methods for details) from test sessions in control (AgRP^tdTomato^) (n=15) and experimental (AgRP^ChR2^) (n=14) mice. Mann-Whitney U Test, U=152, p<0.05. *p<0.05, **p<0.01, ***p<0.001, ****p<0.0001.

### 2.2 Mechanisms underlying the effect of AgRP neuron activity on flavor-nutrient learning

Across all mice, there was a strong correlation between the number of CS+ licks (Figure 2A, R=.636, p<0.001), but not CS- licks (Figure 2B, R=-.159, p=ns), during training and testing. Generally, experimental (AgRP neuron stimulation) mice had a higher number of licks during training (Figure S1B, S1C). Together, these data raised the possibility that AgRP neurons act through either of two non-exclusive mechanisms to affect FNL. First, AgRP neuron activity may increase motivation to consume flavored solutions during training. In this case, increased sampling during training may strengthen the association between the flavors and their respective post-ingestive consequences, thereby increasing CS+ flavor preference. Second, AgRP neuron activity may increase the value of the nutrient upon flavor consumption (indeed, AgRP neuron activity potentiates the dopamine response to food [23]). In this case, AgRP neuron activity may amplify the reward associated with nutrient-paired flavors, such that each lick carries more weight in the animals’ future assessment of the flavors.

**Figure 2.**
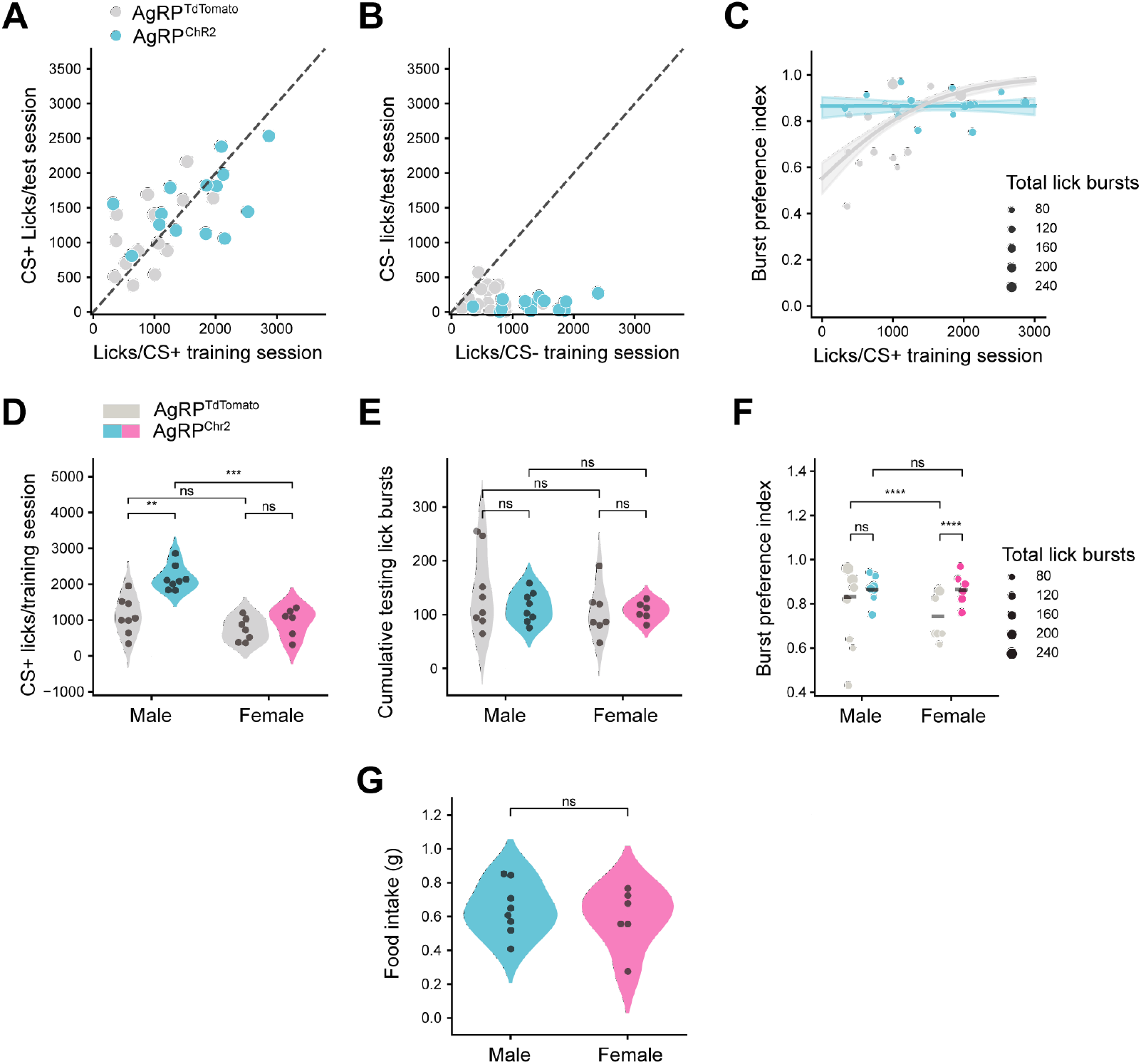
Binomial regression modeling reveals sex differences in the effect of AgRP neuron stimulation on flavor-nutrient learning. (A) Correlation between the average number of CS+ licks during training and testing in control (AgRP^tdTomato^) (n=15) and experimental (AgRP^ChR2^) (n=14) mice. Pearson R, R=.636, p<0.01. (B) Correlation between the average number of CS- licks during training and testing in control (AgRP^tdTomato^) and experimental (AgRP^ChR2^) mice. Pearson R, R=-.224, p=ns. (C) Relationship between average CS+ training licks and burst preference index (from testing) in control (AgRP^tdTomato^) and experimental (AgRP^ChR2^) mice. Cox-Snell R^2^=.980. Shaded region represented 95% confidence interval. (D) Average number of licks per training session during FNL protocol in male and female control (AgRP^tdTomato^) (male: n=8, female: n=7) and experimental (AgRP^ChR2^) (male: n=8, female: n=6) mice. 2-way ANOVA, sex: F(1,25)=27.073, p<0.0001; Chr2: F(1,25)=20.333, p<0.001; sex*Chr2: F(1,25)=7.517, p<0.05. (E) Total number of CS+ and CS- licks in test sessions in male and female control (AgRP^tdTomato^) (male: n=8, female: n=7) and experimental (AgRP^ChR2^) (male: n=8, female: n=6) mice. 2-way ANOVA, sex: F(1,25)=1.701, p=ns; Chr2: F(1,25)=0.694, p=ns; sex*Chr2: F(1,25)=0.711, p=ns. (F) Burst preference index (see Methods for details) from test sessions in male and female control (AgRP^tdTomato^) (male: n=8, female: n=7) and experimental (AgRP^ChR2^) (male: n=8, female: n=6) mice. Lines represent the weighted average within the group, where weights were the corresponding number of lick bursts. Sequential Analysis of Deviance on the Binomial GLM ‘pref ∼ sex*Chr2’ with total lick bursts corresponding to each preference passed as frequency weights, sex: *χ*^2^(1, 27) = 311.492, p<0.001; Chr2: *χ*^2^(1, 26) = 282.295, p<0.0001; sex*Chr2: *χ*^2^(1, 25) = 274.644, p<0.01 AgRP neuron stimulation-evoked food intake (1 h) in male (n=8) and female mice (n=6). T-test, t(12)=-0.577, p=ns. *p<0.05, **p<0.01, ***p<0.001, ****p<0.0001.

To distinguish between these possibilities, we used binomial regression modeling (see Methods for details) to determine whether there was an effect of AgRP neuron stimulation on flavor preference independent of an increase in training licks. Briefly, we modeled flavor preference during FNL testing as a function of the average number of licks per CS+ training session, the AgRP neuron stimulation condition, and their interaction. The model coefficients quantify the effect that these factors have on flavor preference, while the corresponding p-values indicate the probability of the respective coefficients being zero (i.e., the factor having no effect on flavor preference). The resulting model confirmed that mice with more CS+ training licks have significantly stronger preferences during testing (Figure 2C, β=0.0012, t(3430.0)=8.6587, p<0.0001). However, even when accounting for CS+ training licks, there was still a significant, positive effect of AgRP neuron stimulation on preference for the glucose-paired flavor (β=1.6496, t(3430.0)=6.8928, p<0.0001).

We therefore examined interactions between AgRP neuron stimulation and training licks to gain further insight into these effects. The negative coefficient for the interaction between ChR2 expression and CS+ training licks (β=-0.0012, t(3430.0)=-6.8453, p<0.0001) equaled the magnitude of the coefficient for CS+ training licks alone (β=0.0012, t(3430.0)=8.6587, p<0.0001), suggesting that there was no effect of CS+ training licks on preference in mice receiving AgRP neuron stimulation. Indeed, when fitting separate models for control and AgRP neuron-stimulated mice, CS+ training licks were a significant predictor of preference index in control (β=0.0012, t(1874.0)=8.6587, p<0.0001) but not AgRP neuron-stimulated (β=6.31×10^-7^, t(1556.0)=0.006, p=ns) mice. Thus, our models suggest that while CS+ training licks are positively correlated with flavor preferences, the AgRP stimulation-induced increase in CS+ training licks does not fully account for the increase in preference observed in AgRP-stimulated mice. This indicates that, at least for a subset of mice, AgRP neuron stimulation strengthens nutrient-paired flavor preference independently of increasing training licks.

Remarkably, the effects of these two factors on FNL were largely resolved when we analyzed sex differences in the impact of AgRP neuron stimulation on training licks. Specifically, only male mice significantly increased glucose-paired CS+ training licks in response to AgRP neuron stimulation (Figure 2D; male: t(14)=4.804, p<0.01; female: t(11)=1.079, p=ns), even though male and female ChR2-expressing mice had similar preferences for the nutrient-paired flavor (Figure 2E, 2F, β=0.011, t(1556)=0.070, p=ns). In other words, female mice displayed significant increases in CS+ preference with AgRP neuron stimulation (Figure 2F, β=0.790, t(1376)=5.531, p<.0001), despite no increase in training licks (Figure 2D). Conversely, control male mice already displayed high preferences for CS+, such that the increase in preference by AgRP neuron stimulation was not statistically significant (Figure 2F, β=0.267, t(2054)=2.132, p=ns). Further, when fitting a model to data from male mice only, all increases in CS+ preference due to AgRP neuron stimulation were explained by the increase in training licks (ChR2: β=1.320, t(2052.0)=1.951, p= ns; csp_train: β=0.0014, t(2052.0)=7.955, p<0.0001; ChR2:csp_train:β=-.0012, t(2052.0)=-3.4857, p<0.001; ChR2=experimental group, csp_train=CS+ training licks), although it should be noted that there was significant multicollinearity in this model owing to the consistently high number of CS+ training licks in ChR2-expressing male mice. We found the opposite in female mice: the increase in CS+ preference was not mediated by increases in training licks (Chr2: β=1.501, t(1374.0)=3.439, p<0.01; csp_train: β=0.0004, t(1374.0)=1.453, p= ns; Chr2:csp_train:β=-.0008, t(1374.0)=-1.800, p=ns).

Together, these results suggest that different mechanisms underlie the stimulation-induced increase in flavor preference for male and female mice, despite no sex differences in the ability for AgRP neuron stimulation to increase food intake (Figure 2G, t(12)=-0.577, p=ns). In male mice, AgRP neuron stimulation increased consumption during training (and thus the opportunity to learn the flavor-nutrient association), which correlated with higher preferences for the nutrient-paired flavor. In female mice, AgRP neuron stimulation increases flavor-nutrient preferences independently of training licks, suggesting that it enhances the reward value of nutrient-paired flavors. A prediction of this model is that male AgRP neuron-stimulated mice that fail to increase their intake in response to stimulation should *not* have increased preferences. Few ChR2-expressing male mice licked comparably to controls, and therefore we designed a subsequent experiment to directly test this hypothesis.

### 2.3 Preventing AgRP neuron stimulation-induced overconsumption during training blocks flavor-nutrient preferences in male mice

To experimentally test the predictions of our model, we next asked whether AgRP neuron stimulation would still increase flavor preferences when consumption during training is limited. We performed the standard FNL protocol in male and female mice, but limited CS+ licks during training in AgRP neuron-stimulated mice to the average number of licks of the control mice. By design, the number of CS- and CS+ licks during training were equivalent between control and AgRP neuron stimulated mice (Figure 3A, Chr2: F(1,35)=0.262, p=ns; Chr2*CS: F(1,35)=0.830, p=ns). When training licks were restricted, AgRP neuron stimulation had no effect on the total number of licks during testing (Figure 3B, 3C, t(35)=0.045, p=ns) nor on preference index (Figure 3D, U=178, p=ns). Additional analyses revealed sex differences consistent with the previous results. While AgRP neuron stimulation did not affect total licking during testing in either sex (Figure 3E, male: t(16) = −1.895, p=ns; female: t(17) = −2.055, p=ns), female but not male mice with AgRP neuron stimulation displayed significantly stronger preferences for the nutrient-paired flavor than their control counterparts (Figure 3F, male: β=-0.269, z=-2.868, p<.001; female: β=0.349, z=3.444, p<.001). Therefore, our combined data reveal that training licks predict flavor preference in male mice, but AgRP neuron activity enhances the reward value of nutrient-paired flavors independently of training licks in female mice.

**Figure 3.**
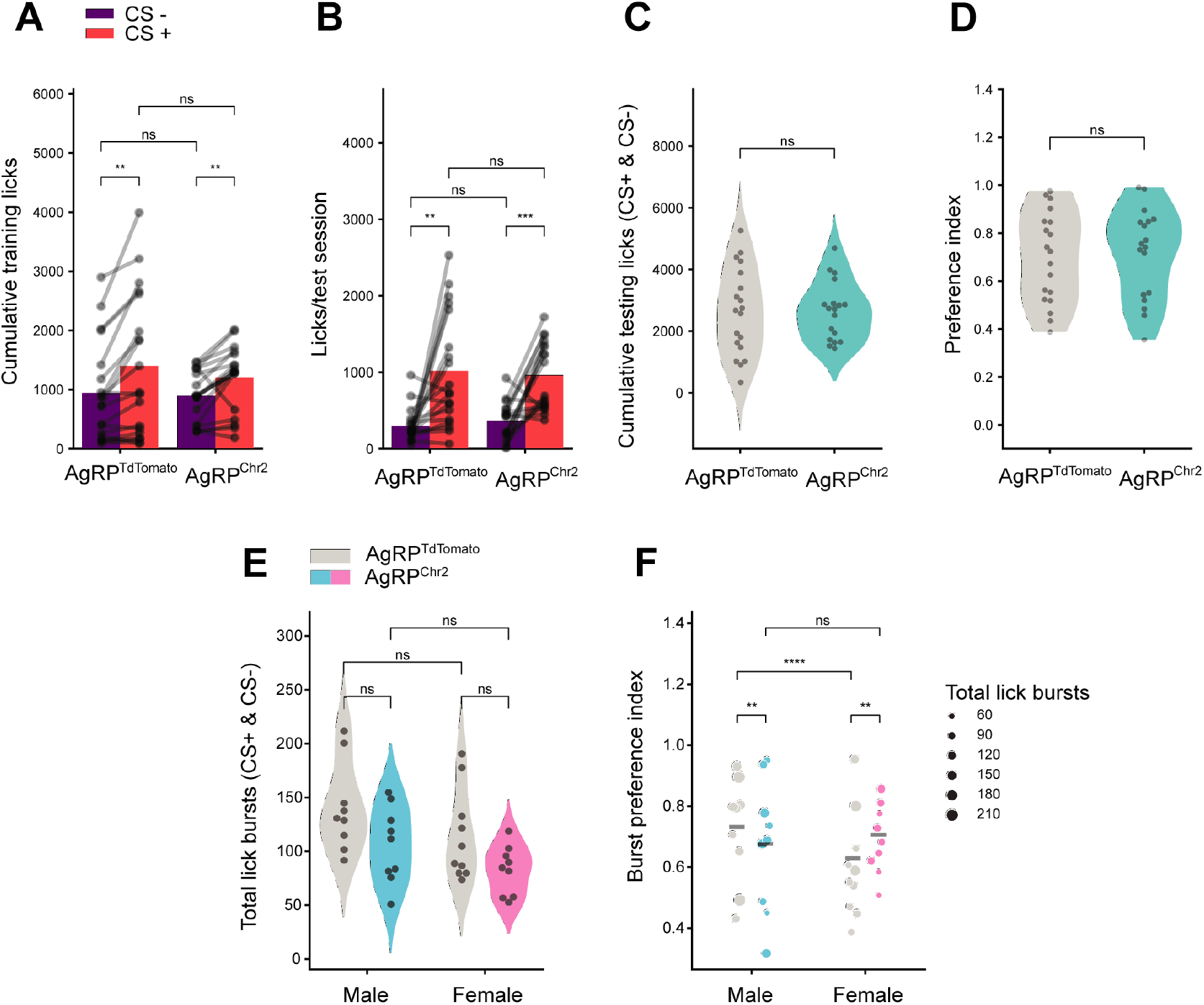
AgRP neuron stimulation increases flavor-nutrient learning independently of training licks in female but not male mice. (A) Average number of licks per training session during limited-intake FNL protocol in control (AgRP^tdTomato^) (n=19) and experimental (AgRP^ChR2^) (n=18) mice. Mixed ANOVA, Chr2: F(1,35)=0.262, p=ns; CS: F(1,35)=30.052, p<0.0001; Chr2*CS: F(1,35)=0.830, p=ns. Bar graphs represent data mean. (B) Average number of licks per test session following limited-intake FNL protocol in control (AgRP^tdTomato^) (n=19) and experimental (AgRP^ChR2^) (n=18) mice. Mixed ANOVA, Chr2: F(1,35)=0.002, p=ns; CS: F(1,35)=32.737, p<0.0001; Chr2*CS: F(1,35)=0.251, p=ns. (C) Total number of CS+ and CS- licks in test sessions in control (AgRP^tdTomato^) (n=19) and experimental (AgRP^ChR2^) (n=18) mice during limited-access FNL protocol. T-test, t(35)=0.045, p=ns. (D) Preference index from test sessions in control (AgRP^tdTomato^) (n=19) and experimental (AgRP^ChR2^) (n=18) mice following limited-access FNL protocol. Mann-Whitney U-test, U=178, p=ns. (E) Total number of CS+ and CS- licks in test sessions in male and female control (AgRP^tdTomato^) (male: n=9, female: n=10) and experimental (AgRP^ChR2^) (male: n=9, female: n=9) mice following limited-access FNL protocol. 2-way ANOVA, sex: F(1,33)=4.480, p<0.05; Chr2: F(1,33)=7.533, p<.01; sex*ChR2: F(1,33)=0.015, p=ns. (F) Burst preference index from test sessions in male and female control (AgRP^tdTomato^) (n=19) and experimental (AgRP^ChR2^) (n=18) mice following limited-access FNL protocol. Lines represent the weighted average within the group, where weights were the corresponding number of lick bursts. Sequential Analysis of Deviance on the Binomial GLM ‘pref ∼ sex*Chr2’ with total lick bursts corresponding to each preference passed as frequency weights, sex: *χ*^2^(1, 35) =582.495, p<.001; Chr2: *χ*^2^(1, 34) = 582.413, p=ns; sex*Chr2: *χ*^2^(1, 33) = 562.258, p<0.0001 *p<0.05, **p<0.01, ***p<0.001, ****p<0.0001.

### 2.4 AgRP neurons track and update associations between flavors and nutrients on short timescales

In the context of eating, AgRP neurons are responsive to sensory cues (acute/transient activity reductions over seconds [4; 18; 19]) and post-ingestive detection of nutrients (long-term/sustained activity reductions over tens of minutes [20; 21; 27]). The data presented thus far address the role of sustained AgRP neuron activity during FNL through optogenetic activation studies. To determine whether and how endogenous AgRP neuron activity tracks FNL across both time scales, we next monitored *in vivo* calcium dynamics of AgRP neurons using fiber photometry. To monitor neural activity in the mouse’s home cage during FNL, we designed custom lickometers (Figure S2A) enabling us to correlate changes in AgRP neuron activity to CS+ and CS- consumption. To validate this equipment, we compared distributions of inter-lick intervals (ILIs) recorded by our custom lickometers to ILIs recorded by commercial Med Associates lickometers and found both systems to be equally precise and accurate at measuring discrete mouse licks (Figure S2B, S2C). We therefore used this system to monitor licking concurrent with AgRP neuron fiber photometry during FNL training and testing sessions.

We injected *Agrp-Ires-Cre* mice with a viral vector encoding the calcium indicator GCaMP6s and implanted an optic fiber above the injection site (Figure 4A, 4B). As previously reported, AgRP neuron activity in food-restricted mice was rapidly suppressed in response to re-feeding (Figure 4C). AgRP neuron activity across training sessions was largely dominated by the long-term, sustained shifts in activity caused by gastric infusions (Figure 4D-G). Specifically, CS+ sessions were accompanied by significant decreases in activity which lasted most of the 30-min session, whereas baseline activity was unchanged throughout CS- sessions. Across training days, the only significant change in the mean response was between training days 1 and 2 (Figure 4E, 4F, Day 1 vs Day2 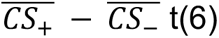 =-6.520, p<0.01), which reflects the flipped ordering of CS+ and CS- sessions on day 2 (for a counterbalanced design, CS+ was provided in the afternoon on days 1 and 3 and on the morning of day 2 of training across experiments; Figure S1A). This is consistent with previous findings demonstrating that AgRP neuron activity changes between morning and afternoon [19]. Overall, the average AgRP neuron activity response across training days was significantly different between CS- and CS+ sessions (Figure 4G, t(6)=-6.881, p<0.001).

**Figure 4.**
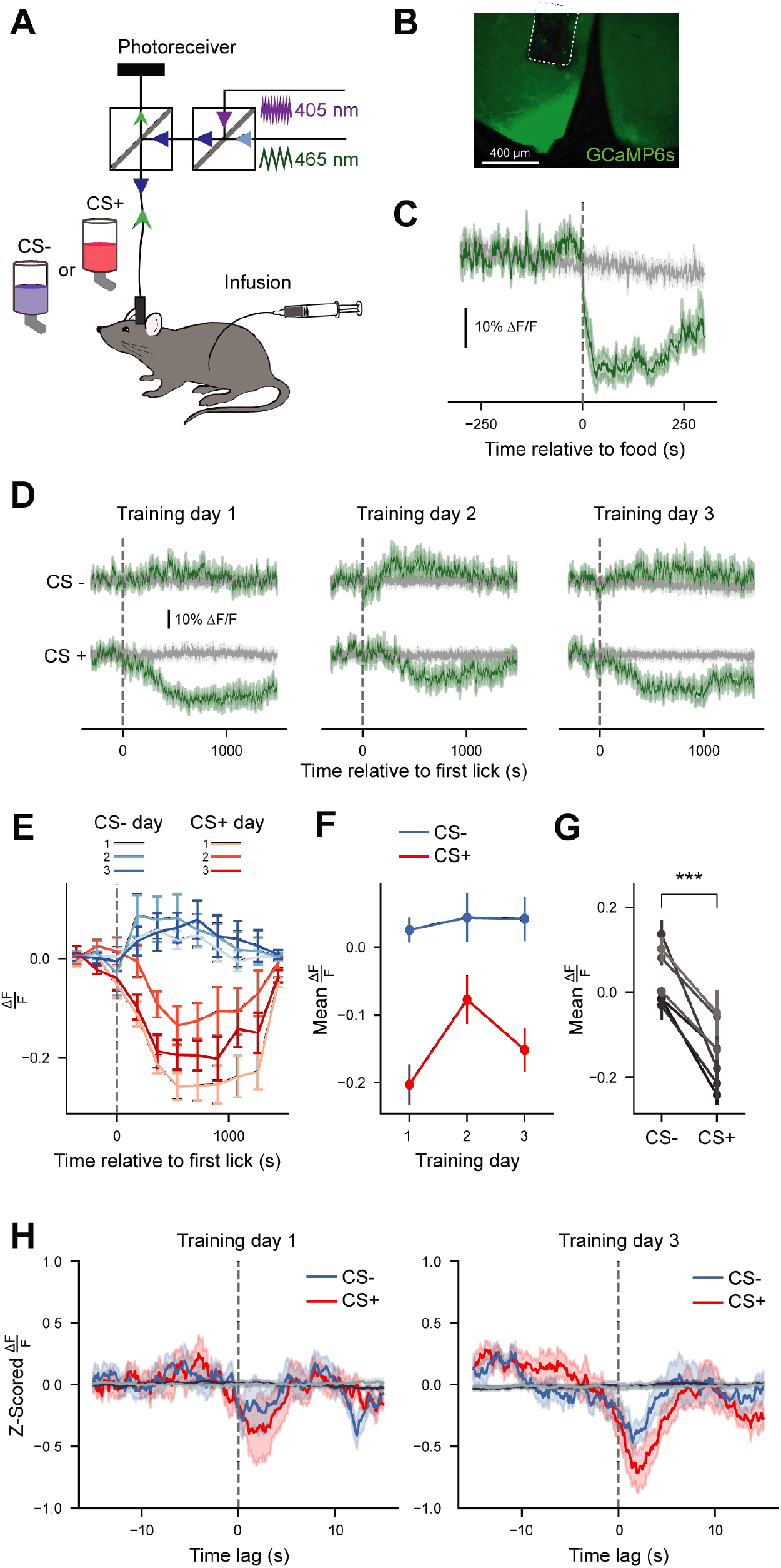
AgRP neurons track associations between sensory cues and nutrients on short timescales during FNL training. (A) Schematic of fiber photometry setup during FNL training sessions. (B) Representative image of GCaMP expression in AgRP neurons of the arcuate nucleus. (C) Average 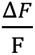 of GCaMP6s signals from AgRP neurons in response to refeeding after an overnight fast (n=7). Green trace, 465-nm calcium-dependent wavelength; gray trace, 405-nm calcium-independent (isosbestic) wavelength. (D) Average 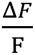 of GCaMP6s signals from AgRP neurons during all training sessions of FNL. Individual traces are aligned to the time of the first lick (n=7). (E) Data from (D), 465-nm signal, binned in 3-min intervals (n=7, see Table S1 for 3-way Repeated Measures ANOVA table). (F) Average mean 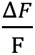 of 465-nm signal from the time of the first lick until the end of each training session across days (n=7). 2-way Repeated Measures ANOVA, day: F(2, 12)=7.353, p<0.05; CS: F(1,6)=47.349, p<0.01; day*CS: F(2,12)=11.104, p<0.01. (G) Average mean 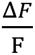 of 465-nm signal from the time of the first lick until the end of sessions averaged across training days. Paired T-test, t(6) = −6.881, p<0.01. (H) Mean bout-triggered average 465-nm responses across training for CS+ (red traces) and CS- (blue traces) sessions. Bout-triggered averages computed from randomly generated lick bouts are shown in dark purple (for CS+) and gray (for CS-). ANOVA on Linear Mixed Effects Model, time*day*CS: F(8, 13922.777)=4.509, p<0.0001 (see Table S1 for full ANOVA table). All data are expressed as mean ± SEM. *p<0.05, **p<0.01, ***p<0.001, ****p<0.0001.

To determine whether there were acute AgRP neuron activity changes over the course of training, we extracted the calcium dynamics surrounding the onset of each lick bout (15 s before and 15 s after lick bout) after normalizing relative to the changing baseline activity (Figure S3, see Methods for details). We then averaged these responses across bouts for each mouse per session to obtain bout-triggered average acute (“peri-lick bout”) AgRP neuron responses (Figure 4H). Given the effect of time of day on AgRP neuron activity, we focused our analyses on training days 1 and 3, when the timing of the training sessions were consistent. This analysis revealed a significant three-way interaction between CS, training day, and time (F(8, 13922.78)=4.509, p<0.0001), reflecting the fact that peri-lick bout CS+ responses are reliably greater in magnitude than peri-bout CS- responses at the end (day 3) of training across both sexes (Figure 4H, day(3):time(5):cs(+): β=-0.598, t(13966.44)=-2.339, p<0.05).

As in previous experiments, after training mice were assessed during two test days where they could choose between licking CS+ or CS- flavors (Figure 5A). Similar to prior results, mice significantly preferred the CS+ over the CS- flavor (Figure 5B, 5C, t(6)=6.857 p<0.001), and tonic AgRP neuron activity was dominated by the (lack of) post-ingestive signaling (Figure 5D). At the time where we observed significant differences between CS+ and CS- peri-lick bout activity during training, responses to CS+ were no longer more negative than CS- (time(5):cs(+): β=0.764, t=2.206, p<0.05). It should be noted, however, that we excluded mice with low numbers (less than 3) of lick bouts to ensure that the bout-triggered average responses were robust and representative – several mice had virtually no intake of the CS- flavor given their strong preference for the CS+ flavor. As a result, the analysis in Figure 5E was underpowered and should therefore be interpreted with caution.

**Figure 5.**
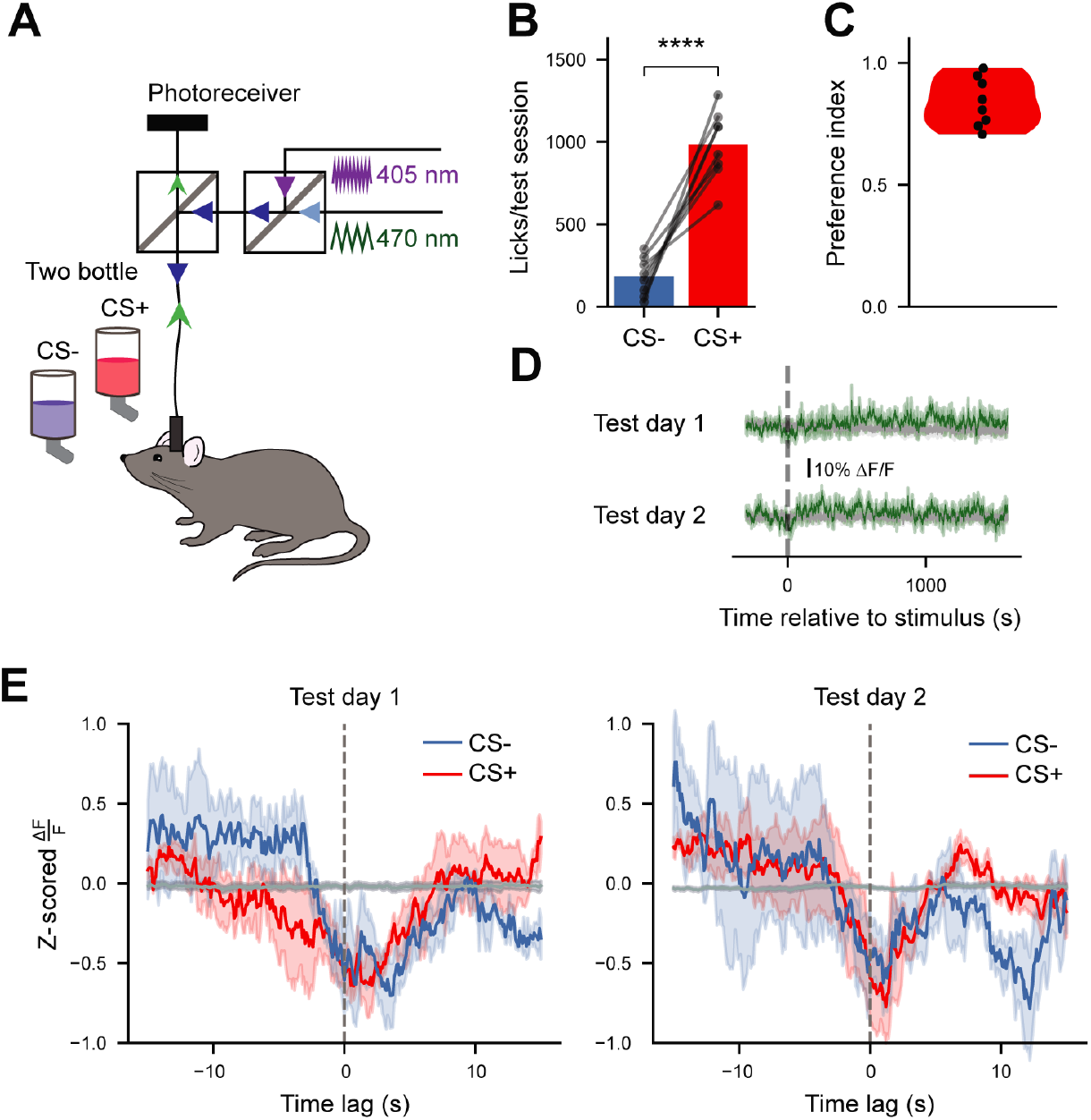
Enhanced acute AgRP responses to CS+ flavors extinguish in the absence of post-ingestive signaling. (A) Schematic of fiber photometry setup during FNL testing sessions. (B) Average CS- and CS+ licks per testing session (n=7). Paired T test, t(6) = 6.857, p<0.01. (C) Preference indices for all mice during testing (n=7). (D) Average 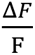 of GCaMP6s signals from AgRP neurons during FNL testing sessions. Individual traces are aligned to the time of the first lick (n=7). Green traces, 465-nm calcium-dependent wavelength; gray traces, 405-nm calcium-independent (isosbestic) wavelength. (E) Mean bout-triggered average responses for both days of testing for CS+ (red traces) and CS- (blue traces) bouts (see Table S1 for ANOVA table) [n=4, mice with too few (< 3) CS- lick bouts during testing were excluded from analyses to ensure that lick bout averages were robust and representative]. Bout-triggered averages computed from randomly generated lick bouts are shown in gray. Data are expressed as mean ± SEM. *p<0.05, **p<0.01, ***p<0.001, ****p<0.0001.

Together, these data indicate that acute, peri-lick bout AgRP neuron activity changes are updated with experience to distinguish between flavors that are associated with nutrients and those that are not. Further, these activity transients extinguish in the absence of post-ingestive (nutrient) feedback.

## 3. Discussion

AgRP neurons are responsive to both sensory and nutritive properties of food [4; 18; 20; 21; 27]. Here, we employed optogenetics and *in vivo* calcium imaging to test how AgRP neuron activity both modulates and responds to FNL. Our results demonstrate that AgRP neuron activity potentiates FNL via sex-dependent mechanisms. In male mice, AgRP neuron activity increased consumption during training which correlated with increased flavor preference. In female mice, the effect of AgRP neuron stimulation on FNL was independent of flavor consumption during training, suggesting that AgRP neuron activity intrinsically increases the reward value of nutrient-paired flavors. We also reveal that AgRP neurons track consumption of these flavors over rapid time scales (i.e. in the seconds surrounding lick bouts), and update activity responses to sensory cues by integrating post-ingestive feedback. Taken together, these data improve our knowledge of the role of central feeding neurons in mediating FNL.

Given that mice prefer flavors associated with AgRP neuron inhibition [4], we initially hypothesized that preventing AgRP neuron inhibition with optogenetic stimulation would block FNL. However, our data demonstrated the opposite: that AgRP neuron activity in hungry mice *potentiates* FNL. In addition to showing that AgRP neuron suppression is not a necessary link in the gut-brain reward pathways for FNL, this suggests AgRP neurons may instead underlie the modulatory effects of energy status on FNL, as both food deprivation [28] and AgRP neuron stimulation (Figures 1-3) enhance FNL (although *ad libitum*-fed rats still exhibit robust FNL [28]). Therefore, in contrast to dopamine signaling which appears essential for FNL [10], AgRP neuron activity has a faciliatory role that amplifies preferences for nutrient-paired flavors. Interestingly, AgRP neuron activity does not influence tonic dopamine signaling but potentiates the dopamine response to food [23]. It is unknown if AgRP neuron activity also potentiates the dopamine response to food-associated cues, such as the CS+ flavors used in the current study.

How does AgRP neuron activity enhance FNL? We used a model-based approach to generate predictions about whether AgRP neuron activity (1) increases flavor consumption during training to facilitate the learned flavor-nutrient association and/or (2) increases the reward value of the nutrient-paired flavor during training. Empirical testing of these predictions revealed that for male mice, AgRP neuron activity primarily increased flavor consumption during training, strengthening the association between flavor and nutrients without increasing the reward value of the nutrient-paired flavor. In contrast, AgRP neuron activation in female mice enhanced flavor preferences independently of increasing consumption, an effect that was consistent across multiple experiments. This result implies that AgRP neuron activity itself, likely via engagement of one or multiple downstream targets [29; 30], increases the intrinsic reward value of the nutrient-paired flavor. To our knowledge, this is the first report of AgRP neuron stimulation causing sexually dimorphic feeding behavior, although an interesting recent study demonstrated sex differences in how stress changes preference for AgRP neuron stimulation: a subset female, but not male, mice prefer rather than avoid AgRP neuron stimulation after stress [31]. Further, there are known sex differences regarding the function of metabolic signaling pathways in AgRP neurons [32; 33] and Agrp protein expression [34–36]. Because AgRP neuron stimulation increases food intake equally in males and females (Figure 2G), it will be interesting to investigate the mechanisms that underlie the observed sex differences specifically related to FNL. For example, it is possible that there are differences between males and females in sensory processing, gut-brain signaling, or associative learning that may underlie these differences.

In these studies, we used binomial regression modeling to analyze our FNL data for a better understanding of how AgRP neuron activation increases preference for nutrient-paired flavors. While binomial regression modeling is a well-established technique that has been applied to choice data in behavioral neuroscience applications [37; 38], this is the first instance of its use with data pertaining to FNL. We suggest that FNL is a natural use case for this type of modeling, as it offer researchers the ability to generate and test additional hypotheses about the factors contributing to the development of flavor preferences. In our case, these models enabled us to clarify the effect of a mouse’s experience with the CS+ flavor during training on their preference during testing, and revealed underlying sex differences.

Across our behavioral studies, we performed optogenetic stimulation in food-restricted mice to prevent the AgRP neuron inhibition that would normally occur in response to nutrients (Figure 1A). A limitation of this approach, however, is that it is unclear to what extent this optogenetic manipulation prevents neural activity reduction versus artificially stimulates AgRP neurons beyond normal levels in hunger. To address this concern, we stimulated AgRP neurons at 20 Hz, which is the frequency of AgRP neuron firing in food-restricted mice [19]. Nonetheless, the supra-physiological nature of optogenetic stimulation limits interpretations of these results.

In addition to determining how exogenous AgRP neuron activity influences FNL, we recorded endogenous activity in AgRP neurons to determine natural activity patterns during the development of associations between flavors and nutrients. Because AgRP neurons respond to sensory cues that predict nutrients [4; 18; 19], we were somewhat surprised that we did not observe changes in tonic AgRP neuron activity in response to CS+ vs. CS- across training or testing. However, this finding is consistent with previous work demonstrating that long-term AgRP neuron activity levels are dominated by post-ingestive signaling [21]. Differences in AgRP neuron responses to CS+ and CS- were instead observed when we analyzed acute AgRP neuron activity responses to individual lick bouts during training. Indeed, AgRP neurons discriminate between CS+ and CS- across training days. Importantly, these activity changes do not simply reflect the presence of nutrients in the gut: because the intra-gastric glucose infusion is identical on training days 1 and 3, the effect of the infusion on AgRP neuron activity is expected to be similar. Nonetheless, by the end of training, licking for the CS+ flavor produced a more robust AgRP neuron activity transient than at the beginning of training. These differential activity dynamics to CS+ and CS- flavors suggest that acute, peri-bout AgRP neuron activity reflects the learned association between flavor and nutrient. In other words, the acute changes in AgRP neuron activity track a prediction of nutrition over the course of training. Therefore, it makes sense that these acute AgRP neuron activity changes rapidly extinguish during testing, in the absence of post-ingestive signaling.

Our peri-bout AgRP neuron activity analyses complement *in vivo* electrophysiological [19] and fiber photometry [18] data demonstrating that AgRP neuron activity is modulated by licking for a liquid diet over fast (seconds) timescales. Interestingly, the electrophysiological data revealed considerable variability in individual AgRP neuron responses: about half were inhibited and half were activated by licking (although few AgRP neurons that responded to licking were recorded) [19]. Our fiber photometry data indicate that, at the population level, AgRP neuron activity is generally suppressed by flavor consumption, effects that are significantly enhanced with licking for nutrient-paired versus unpaired flavors. It will be important to perform similar studies with single cell resolution to determine heterogeneity in AgRP neuron responses to nutrient-paired flavors. Together with this previous work, our findings raise the possibility that smaller, acute AgRP neuron activity changes are physiologically relevant for feeding behavior and/or for learning flavor-nutrient associations. Therefore, it will be interesting for future experiments to trigger acute AgRP neuron stimulation/inhibition during lick bouts to determine the behavioral consequences of these activity transients. Finally, it is worth noting that our evaluation of AgRP neuron activity did not reveal sex differences, but given that this particular analysis was underpowered, this could be revisited and confirmed in future studies. Overall, our data add to the literature showing that nutrients train AgRP neurons to predict the nutritive value of food via sensory properties [21], and importantly, provide insight into how AgRP neurons track this information on short timescales.

## 4. Conclusions

FNL is fundamental to survival – across species, animals and humans alike use sensory cues to learn about foods that are nutritious and those that are not (or, that are potentially dangerous). Our work demonstrates that AgRP neuron activity enhances the expression of flavor-nutrient preferences and highlights a role for AgRP neurons in the rapid tracking of flavor-nutrient associations. Overall, this work provides valuable insight into the neural mechanisms underlying FNL.

## Abbreviations

AgRP: agouti-related protein
ChR2: channelrhodopsin-2
CS: conditioned stimulus
FNL: flavor-nutrient learning
ILIs: inter-lick intervals

## Acknowledgements

We thank M. Dietrich for comments on the manuscript and A. Acosta, B. Greenwald, and M. Smith for technical assistance. A.L.A. is a New York Stem Cell Foundation – Robertson Investigator and a Pew Biomedical Scholar. This work was supported by the National Institutes of Health [R00DK119574, R01DK131558 and DP2AT011965 to A.L.A.; N.T.N. (3R00DK119574-05S1) and A.V. (3DP2AT001965-01S1) were supported by NIH Research Supplements to Promote Diversity in Health-Related Sciences], the American Heart Association (857082 to A.L.A.), the New York Stem Cell Foundation (to A.L.A.), the Klingenstein Fund and Simons Foundation (to A.L.A.), the Pew Charitable Trusts (to A.L.A.), and the Monell Chemical Senses Center (to A.L.A.).

## Data and code availability

All code used in this study can be accessed at https://github.com/nathanielnyema/fnc_agrp_project and https://github.com/nathanielnyema/lickometer. All data will be provided on figshare upon publication.

## Author contributions

**Nathaniel T. Nyema:** Conceptualization, methodology, software, validation, formal analysis, investigation, data curation, writing – original draft, visualization, project administration. **Aaron D. McKnight:** Conceptualization, methodology, investigation, writing – review and editing. **Alexandra G. Vargas-Elvira:** Methodology, investigation, writing – review and editing. **Heather M. Schneps:** Investigation, writing – review and editing. **Elizabeth G. Gold:** Investigation. **Kevin P. Myers:** Methodology, writing – review and editing. **Amber L. Alhadeff:** Conceptualization, methodology, resources, writing – original draft, visualization, supervision, project administration, funding acquisition.

## 5. Material and methods

### 5.1 Subjects

Mice were singly housed on a 12-h light/12-h dark cycle with *ad libitum* access to food (Purina Rodent Chow, 5001) and water except under conditions of food restriction or when otherwise noted. All mice were at least 8 weeks old prior to experimentation. *Agrp-Ires-Cre* (Agrptm1(cre)Lowl/J) [39], *Ai32* (B6;129S-Gt(ROSA)26Sortm32(CAG-COP4*H134R/EYFP)Hze/J) [40], and *Ai9* (B6.Cg-Gt(ROSA)26Sortm9(CAG-tdTomato)Hze/J) [40] mice crossed to a *C57BL/6J* background were used. Experiments were performed in both male and female subjects. All procedures were approved by the Monell Chemical Senses Center Institutional Animal Care and Use Committee.

### 5.2 Experimental Procedures

#### 5.2.1 Fiber optic implantation and viral injection

Mice were anesthetized in an induction chamber with 1.5%–3% isoflurane (Clipper, 0010250) and placed in a stereotaxic apparatus. Viral injections were performed as previously described [41]. For somatic stimulation of AgRP neurons, *Agrp-Ires-Cre* mice were crossed to an *Ai32* reporter line to express Channelrhodopsin-2 (ChR2) in AgRP neurons (*Agrp-Ires-Cre*::*Ai32* mice). For control mice *Agrp-Ires-Cre* mice were crossed to an Ai9 reporter line to express tdTomato in AgRP neurons (*Agrp-Ires-Cre::Ai9* mice). A custom-fabricated ferrule capped optical fiber (ferrule: Kientech, FZI-LC-230; fiber: 200-µm core, NA 0.37, ThorLabs, FT200UMT) was then placed unilaterally over the arcuate hypothalamic nucleus (ARC; 1.35 mm posterior to bregma, 0.23 mm lateral to midline, 5.95 mm ventral to the skull) in both *Agrp-Ires-Cre*::*Ai32* and *Agrp-Ires-Cre*::*Ai9* mice and cemented to the skull with Metabond cement (Parkell, S380) and dental cement (Lang Dental Manufacturing, Ortho-jet BCA Liquid, B1306 and Jet Tooth Shade Powder, 143069). For fiber photometry, unilateral injections of a virus designed to Cre-dependently express GCaMP6s (AAV1.Syn.Flex.GCaMP6s.WPRE.SV40, Addgene, 100845-AAV1) were performed in the arcuate nucleus of the hypothalamus (ARC) (300 µL total) in *Agrp-Ires-Cre* mice (1.35 mm posterior to bregma, 0.23 mm lateral to midline, 6.15-6.3 mm ventral to the skull). A ferrule-capped optical fiber (400 µm core, NA 0.48, Doric, MF2.5, 400/430-0.48) was then implanted 0.2 mm above the injection site and secured to the skull with cement as described above.

#### 5.2.2 Gastric catheter implantation

Gastric catheters were assembled by placing surgical mesh (5-mm diameter piece, Bard, 0112660) and an epoxy ball on the end of a 7-cm segment of Micro-Renathane catheter tubing (Braintree Scientific, MRE-033). The tubing was coupled to another segment of the same tubing with an L-shaped 26-g connector secured to another piece of surgical mesh. Mice were anesthetized with isoflurane (1.5%–3%) and an abdominal midline incision was made through the skin and muscle. The end of the catheter with the epoxy ball was then inserted into the fundus of the stomach through a puncture hole and secured in place with the surgical mesh. The other end of the catheter was routed through an intrascapular incision and secured in place around the L-shaped junction with sutures. The catheter was flushed with sterile water and sealed with a metal cap to prevent blockage. Mice were fed with moistened chow and body weight was monitored until pre-surgical weight was regained before starting experiments.

#### 5.2.3 *In vivo* photostimulation

Optogenetic stimulation of AgRP neurons was performed as we have previously described [41]. Briefly, an Arduino UNO was programmed to generate TTL pulse trains with 10-ms pulses firing at 20 Hz with a 1 s ON time followed by a 3 s OFF time. The TTL pulses from the Arduino controlled a 1W 450-nm laser (Lasever, LSR450NL-1W-FC+LSR-PS-II) which was coupled to a multimode optical fiber (200-µm core, NA 0.37, Doric, MFP_200/220//900-0.37_2m_FCM_FCM) with a 1.25-mm OD zirconia ferrule and mating sleeve (Kientech, FZI-LC-230) via a single channel fiber-optic rotary joint (Doric, FRJ_1x1_FC-FC). The power output at the tip of the optical fiber was set to approximately 30 mW before all sessions.

#### 5.2.4 Verification of AgRP neuron stimulation

To functionally confirm optical fiber placement above the ARC, mice were screened for increased food intake during optogenetic AgRP neuron stimulation as we and others have reported previously [2; 41]. For at least one hour, mice were habituated to a chamber with a lined floor and ad libitum access to chow and water. After habituation, food intake in the absence of photostimulation was measured for 1 h to establish baseline intake.

Photostimulation was performed during the following hour. Mice that consumed < 0.3 g of chow during photostimulation were excluded from subsequent experiments.

#### 5.2.5 Food restriction

For experiments performed in food restriction, mice were maintained at ∼90% of baseline body weight. Each mouse was given a ration of 2-3 g of chow at least 1 h after their final training or testing session for the given day.

#### 5.2.6 Flavor-nutrient learning experimental procedures

##### Apparatus

Training sessions were performed in 8 identical Med Associates operant conditioning chambers housed separately within sound-attenuating boxes. Each chamber was equipped with a 1W laser driven by an Arduino UNO, a computer-controlled syringe pump (Med Associates, PHM-210), infusion tubing and wire grid flooring (Med Associates, ENV-307W-GFW). For all habituation and training sessions, a single bottle holder was placed in the center of the wall of the apparatus to avoid the development of a side preference. The walls of the chamber were rearranged prior to two-bottle choice testing to accommodate two bottle holders, one on either side of the wall. The floor and bottle holders were all wired to a contact lickometer (Med Associates ENV-350CW and ENV-250). Custom MED-PC code recorded the time offsets of lick events throughout all sessions and triggered a gastric infusion upon the 20^th^ lick during late habituation sessions and all training sessions. This gastric infusion lasted 10 min and occurred at a rate of 0.06 ml/min for experiments where mice had *ad libitum* access to the flavored solutions. For experiments where mice that had limited access to the flavored solutions infusions were 5 min at 0.12 ml/min to avoid mice receiving infusions without access to the flavored solutions (as we anticipated needing to remove bottles approximately 7 mins after the start of the infusion on average for experimental mice; see Training Procedures for more detail).

##### Training Procedures

A schematic describing training and testing procedures is provided in Figure S1. All mice were habituated to the conditioning chambers for 4 consecutive days prior to the first day of training. Mice were water restricted overnight prior to each of the first 2 days of habituation to motivate licking behavior. During these first 2 sessions, mice were given 0.05% sodium saccharin to consume and received no gastric infusions. Mice were returned to their home cages following each session and given access to water. Mice were subjected to an overnight fast prior to the third day of habituation and maintained at ∼90% of their baseline body weight for the remainder of the experiment as described above. On the last 2 days of habituation mice received an IG infusion of water upon consuming the saccharin solution to habituate to gastric infusions.

Immediately after habituation, mice were exposed to 3 days of training with 2 sessions per day separated by a washout period of at least 3 h. During training sessions, mice had access to 0.05% saccharin flavored with 0.05% Kool-Aid powder (grape and cherry, Amazon). On each training day, mice received both a conditioned stimulus (CS)+ (glucose, 16.67%) and a CS- (water) session in counterbalanced order over the course of training. Grape and cherry flavor pairings were counterbalanced to gastric infusions of glucose or water. Mice were connected to lasers for all training sessions and received AgRP neuron or control photostimulation as described above.

For experiments where intake was limited, bottles containing the flavored solutions were manually removed from the individual apparatus when the mouse’s intake reached a threshold as signaled through MED-PC. To avoid the confound of only experimental mice having their bottles removed during training, we set thresholds for both control and experimental (AgRP neuron stimulation) mice as follows. Control mice were subjected to a threshold on the amount of time they were allowed access to the flavors after the start of infusions based on the expected amount of time it would take experimental mice to reach the lick threshold for a given session. This time threshold was computed using data from the training sessions from previous experiments (i.e., Figures 1 and 2) where mice had *ad libitum* access to the flavored solutions. Specifically, for each training session of the *ad libitum* experiments, we computed a hypothetical lick threshold for the experimental mice by taking the mean number of licks in the control cohort. We then determined for each experimental mouse how much time had passed from the start of their infusion until they reached the session’s hypothetical lick threshold. If a mouse did not reach threshold for a given session, we counted their time until threshold as the duration of the session (30 mins). The average time until threshold for a given session was the amount of time the control mice in the limited intake experiment were allowed access to their bottles from the start of their infusion for the equivalent session. Experimental mice were then subjected to a threshold on the number of licks they were allowed based on the average number of licks that the controls reached for the same session. This strategy successfully enabled us to control for the number of licks between control and experimental mice (Figure 3A), while also controlling for manual interruption during training sessions across groups.

##### Testing procedures

Mice were habituated to the 2-bottle testing environment the day after the last training day. During 2-bottle habituation, mice were presented with one bottle containing 0.05% saccharin and another containing 0.025% saccharin for 30 min. Later in the day, the mice were given these same solutions with the bottle sides flipped. Data were reviewed at the end of the day to ensure all mice sampled both bottles and that there were no side preferences. All mice that met these criteria were then subjected to two 30-min 2-bottle testing sessions with the flavored solutions over two days. The side of the CS+ flavor was counterbalanced across the two days of testing. No photostimulation was performed during testing.

##### Behavioral data analysis

All data analyses were performed using custom Python (3.11.0) scripts. Testing performance was quantified by either a lick or lick-burst based preference index. In both cases, the following formula was used to compute a cumulative preference index for each mouse:

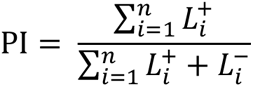

Where *n* is the number of testing days, 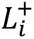 is the number of CS+ licks or lick bursts on day *i* and 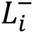 is the number of CS- licks or lick bursts on day *i*. Lick bursts are defined as a sequence of licks of the same flavor where each inter-lick interval is no more than 500 ms.

##### Behavioral Modeling

Preference indices are proportions and are therefore bounded on the interval [0,1]. As a result, these data are poorly characterized by simple linear regression models for two key reasons: (1) they are not meaningfully described by a linear function of any variable considering a line or hyperplane with non-zero slope would imply an unbounded output such that on some domain the model predictions would be invalid (i.e. >1 or <0); and (2) a normal distribution cannot capture the expected asymmetry in the model residuals about predicted preference indices near 0 or 1, where ideally the model should inherently acknowledge that values <0 or >1 have 0 probability. In special cases where the range of proportions to be modeled is small enough and the proportions themselves are close to 0.5, these considerations are less of an issue because within the domain of the data, a linear model is a good enough approximation. In the case of flavor nutrient learning; however, the preferences observed tend to saturate near 1, especially with AgRP neuron stimulation.

A more natural framework for modeling these data is binomial regression, wherein each lick burst can be formalized as a Bernoulli trial, such that a success is a CS+ lick burst and a failure is a CS- lick burst. The number of CS+ lick bursts out of *n* total lick bursts can thus be described as a binomial distributed random variable parametrized by *n* and the expected success rate *p* (preference), which can further be modeled as a function of several independent variables. More formally, for a given mouse:

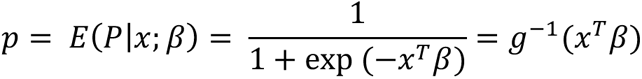

where *x* is a vector of independent variables and *β* is the vector of model coefficients for each variable. In the language of generalized linear models (GLMs), *g*^-1^ is the inverse link function such that 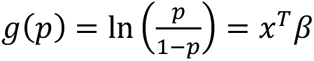 is the link function itself and relates the expected value of the preferences to the linear predictor *x*^*T*^*β*. In other words, the link function is a transformation that linearizes preference indices, in part by converting them to an unbounded quantity. The resulting quantity can be described as a linear combination of certain independent variables of interest via the model coefficients. Importantly, the link function in this case is the logarithm of the odds ratio. As a result, any coefficients for these models should be interpreted in units of log odds.

Importantly, the flavor preference index we use for these models is the proportion of CS+ lick *bursts* as opposed to total CS+ *licks* (the more commonly used parameter). The reason for this is that individual lick events are inadvertently autocorrelated: licks that occur within the same lick burst by definition always correspond to the same flavor. As a result, if we were to define each lick as a trial, the number of successes (CS+ licks) would not follow a binomial distribution which assumes all trials are independent. In practice, this would result in overdispersion [42], a phenomenon where there is greater variability in the data set than predicted by the model. Ultimately, this produces artificially small p-values. Although the lick burst preference index underestimates the lick-based preference index (Signed Rank Test: W=7, p<0.001), these metrics are nonetheless strongly correlated (Figure S4, Spearman Rank Correlation: ρ=0.907, p<0.001). Therefore, this modified preference index is a reasonable and more conservative approximation of the traditionally reported statistic (total licks) for use in modeling-based approaches.

When evaluating the effects of mean CS+ training licks on preference index the following model was used:

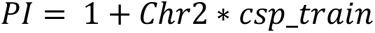

Where Chr2 is dummy coded to represent whether a mouse expressed Channelrhodopsin-2 for AgRP neuron stimulation, and csp_train is the mean number of CS+ training licks. When fitting this model separately for AgRP stimulated and control mice the following simplified model was used instead:

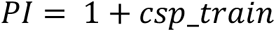

In place of t-tests, when performing pairwise tests of burst preference indices (as shown in Figures 2C, 2F), the following model was fit:

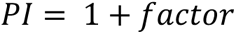

Where *factor* is a dummy coded variable representing participation in one of the two groups to be compared, and the p-value for the coefficient on *factor* is reported in the plots (i.e. Figure 2F & 3F).

All binomial regression models presented in our results were fit with the statsmodels library (0.13.5) in Python (3.11.0) using the statsmodels.formula.api.glm function and passing the total lick bursts corresponding to each mouse’s preference index to the *freq*_*weig*ℎ*ts* argument.

#### 5.2.7 *In vivo* fiber photometry

Fiber photometry was performed as we have previously described [21; 27; 41]. Briefly, GCaMP6s was excited by 465-nm light modulated at 211 Hz while an isosbestic 405-nm light modulated at 566 Hz was used to control for artifacts caused by movement and bleaching. Output power of 465-nm excitation light was adjusted to 50% (20–60 µW) and 405-nm excitation light was adjusted to 5% (2–10 µW) detection range of the photoreceiver to avoid signal saturation. Small power adjustments were made to compensate for variations in fiber placement and viral expression such that all recordings began with approximately the same baseline fluorescence values across all mice. 5-10 min of baseline GCaMP6s fluorescence was measured prior to the beginning of each FNL training and testing session. All data were collected through Synapse Tucker-Davis Technologies software.

#### 5.2.8 Home cage lickometer

To conduct FNL training and testing procedures in a mouse’s home cage during fiber photometry recordings, we designed an Arduino-based home cage lickometer as shown in Figure S2. This device records licks and triggers the appropriate infusion similar to the Med Associates setup. The mechanism of lick detection is similar to that described in other systems [43–45]. The lickometer circuit itself is a resistive voltage divider where the mouse functions as a switch that allows current to flow through the circuit when closed. The mouse remains in contact with the 5V port on the Arduino UNO via an alligator clip attached to wire flooring placed on the bottom of a standard mouse cage. The spout is in turn wired to one terminal of a 3.3 MOhm resistor which is grounded on the other terminal. When the mouse closes the circuit by making contact with the spout, the voltage drop across the resistor increases from 0 V to approximately 2.6 V (assuming the average mouse has an equivalent resistance of approximately 3 MOhms [45]). The voltage drop across the load resistor is read continuously by an analog pin on the Arduino and a buffer of the 6 most recent readings is stored. After each new reading, a de-noised estimate of the change in voltage is computed by averaging the first 3 values in this buffer and the last 3 values separately and taking the difference. A lick is registered when the difference between the current and previous voltage crosses a user-defined threshold which is set separately for all devices. This methodology allows us to programmatically debounce the input signal. Every time a lick is registered, the time is printed to the serial port of the Arduino. A custom Python script which is run during the experiment asynchronously reads this data stream, timestamps the events, and saves the results to a csv file for post-hoc analyses.

This same device was also programmed to trigger the appropriate infusion upon the 20th lick. For the accompanying hardware, a 3-pin Molex adapter is used to gain access to the input pins of a Med Associates syringe pump. These pumps are designed such that the pump is on when the “Operate” pin is grounded. As a result, a MOSFET is used to connect the “Operate” pin to ground whenever a digital pin on the Arduino is programmed to be in the HIGH state (Figure S2A).

#### 5.2.9 Fiber photometry data analysis

##### Pre-processing

Fiber photometry data were analyzed similarly to our previous reports [21; 23; 27; 41] and modified to monitor calcium signaling dynamics around lick bouts. Mice were selected for inclusion in the following analyses based on AgRP dynamics in response to a chow food drop after an overnight fast. Specifically, data from these sessions were first normalized using the following equation:

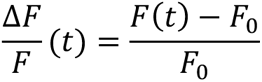

Where *F*_0_ is the median of the raw fluorescence data from either the 465-nm or 405-nm channel in the 5-min pre-stimulus period. Only mice with minimum 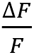 values less than −0.2 (−20%) were included in subsequent analyses.

For all recordings performed during FNL training and testing, the traces from the 465-nm channel were corrected for movement and bleaching artifacts by subtracting calcium-independent fluorescence changes as approximated by a linear transformation of the smoothed 405-nm channel data. Specifically, the 405- and 465-nm channel data were separately downsampled to 1Hz and smoothed by convolving each signal with a 10-sample standard deviation Gaussian kernel. Least squares linear regression was then used to estimate a linear mapping between smoothed 405- and 465-nm channel data. The median of the fitted 405-nm channel data during the 5-minute pre-stimulus period was then subtracted from the entire signal before re-upsampling and subtracting it from the raw 465-nm channel data. Importantly, a non-negativity constraint is placed on the slope in the above regression analysis to avoid over-correcting for strong negative correlations between the 405 and 465 which manifest as positive deflections in the 405-nm signal time-locked to sharp negative deflections in the 465-nm signal (presumably because 405 nm is not the absolutely precise isosbestic point for GCaMP). The detrended 465-nm signal is then downsampled to 10 Hz for computational efficiency and normalized using the following equation:

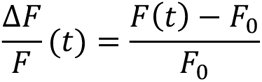

For all analyses of long timescale AgRP activity, *F*_0_ is the median of the corrected signal in the 5-min pre-stimulus period. For all peri-bout analyses, *F*_0_ is instead treated as a function of time representing the estimated baseline activity at time *t*. As such, 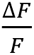 represents a fold change in fluorescence relative to a changing baseline. This baseline is approximated by a moving average of window size 60 s applied to the corrected fluorescence signal (Figure S3) such that:

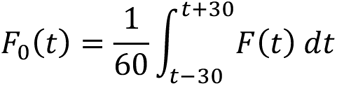

Fluorescence and lick bout data in the first and last 30 s is discarded due to the lack of a complete set of points for baseline calculation. No bouts are discarded in the first 30 s since this is before spout presentation, and there was only ever at most 1 bout in the last 30 s of any session. Finally, for all peri-bout analyses, the 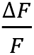 values are further z-scored relative to the mean and standard deviation of the remaining 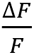 values in the 5-min baseline period prior to spout presentation.

##### Peri-bout analyses

Custom Python scripts were used to map all lick events for each session to the corresponding time stamps in the fiber photometry recordings. The onsets of lick bouts were defined as the time offsets of any licks that were preceded by an inter-lick interval of at least 20 s and followed by at least 1 more lick within less than 20 s. Importantly, for 2-bottle testing, licks were pooled across flavors before computing inter-lick intervals to avoid extracting lick bouts where the animal had licked the opposite flavor less than 20 s prior. The normalized photometry signal 15 s before and after the onset of each bout was extracted to yield a set of peri-bout responses for each recording session. Bout-triggered average AgRP responses were obtained for each recording session by averaging z-scored 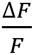 values across bouts for each time lag relative to the respective bout onsets. As a control, for each session for each mouse, random lick bouts were generated and activity around these bouts was extracted and used to compute a randomized bout-triggered average response. This procedure was repeated 100 times for each mouse for each session, and the responses were further averaged across repeats. For each repeat, the number of random lick bouts generated equaled the number of lick bouts achieved by that mouse during that session. The resulting traces were compared to bout-triggered averages obtained from actual lick bouts to verify our observations could not be produced by chance. Additional exclusion criteria were used for peri-bout analyses during testing to ensure the robustness of bout-triggered averages. Specifically, only mice with at least 3 lick bouts on both flavors for both testing days were included in the analysis of testing data.

Linear mixed effects models were used to evaluate the effects of CS, time, and training/testing day peri-bout responses. Individual peri-bout responses are transient and vary in the timing of the peak response. As a result, models that treat time as a categorical variable (i.e. independent coefficients are fit for each time bin) poorly characterize the data. We instead model time with a basis set of 8 degree-3 b-splines (Figure S5), similar to [46]. Conceptually, by fitting the model we are estimating weights for these splines such that their weighted sum reconstructs the mean response for each CS for each day. The full model equation was as follows:

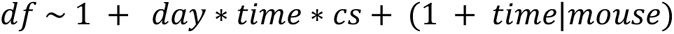

When model coefficients involving the variable *time* are referenced in the text, the number following the variable name indicates the index of the spline that the coefficient is associated with (i.e. time(5) would refer to the fifth spline shown in Figure S5). Data from each individual lick bout across mice were fed into these models. The lmerTest (3.1-3) package in R is used to compute ANOVA tables from the results, using Satterthwaithe’s approximation to derive effective degrees of freedom.

#### 5.2.10 Fos immunohistochemistry

Following all experiments, mice were placed in a cage with only water (no food) and given AgRP neuron (or control) stimulation for 1 h. After 90 min, mice were deeply anesthetized with isoflurane and transcardially perfused with 0.1 M Dulbecco’s phosphate-buffered saline (PBS, HyClone, SH30013.04) followed by 4% paraformaldehyde (MP Biomedicals, 150146). Brains were post-fixed overnight in 4% paraformaldehyde then transferred to 30% sucrose in PBS. 100-µm sections were collected on a cryostat. Sections were incubated in primary cFos antibody [c-fos (9F6) rabbit mAb, Cell Signaling Technology, 2250) at 1:1000 dilution overnight at 4°C. Sections were then washed 3x for 5 min each in PBS at room temperature. Secondary antibodies were applied (Alexa647 donkey anti-rabbit, 711-605-152 or Cy3 donkey anti-rabbit, 711-165-152; Jackson ImmunoResearch. Sections were mounted to slides, coverslipped with Fluorogel, and images were taken on a Leica STELLARIS5 confocal microscope (purchased with S10OD030354, PI: Alhadeff).

#### 5.2.11 Statistical Analyses

All statistical analyses were performed using custom Python (3.11.0) scripts using appropriate functions from the following libraries: Scipy (1.10.1), Pingouin (0.5.3), and statsmodels (0.13.5). For certain analyses where there were no available Python implementations, rpy2 (3.5.7) was used to invoke appropriate functions from lme4 (1.1-31) or lmerTest (3.1-3). For ANOVA models fit to smaller sets of data, the Shapiro-Wilkes test was used to verify normality. For models fit to larger datasets, QQ-plots were inspected instead. Where appropriate, Levene’s test was used to test for equality of variances. Mauchly’s test was used to test for sphericity and when necessary, p-values were Greenhouse-Geisser corrected. For all post-hoc pairwise tests, p-values were adjusted using the Holm-Sidak method. All bar graphs represent data mean. *p<0.05, **p<0.01, ***p<0.001, ****p<0.0001. All statistical information is provided in Tables S1 and S2.

---

**Figure S1.**
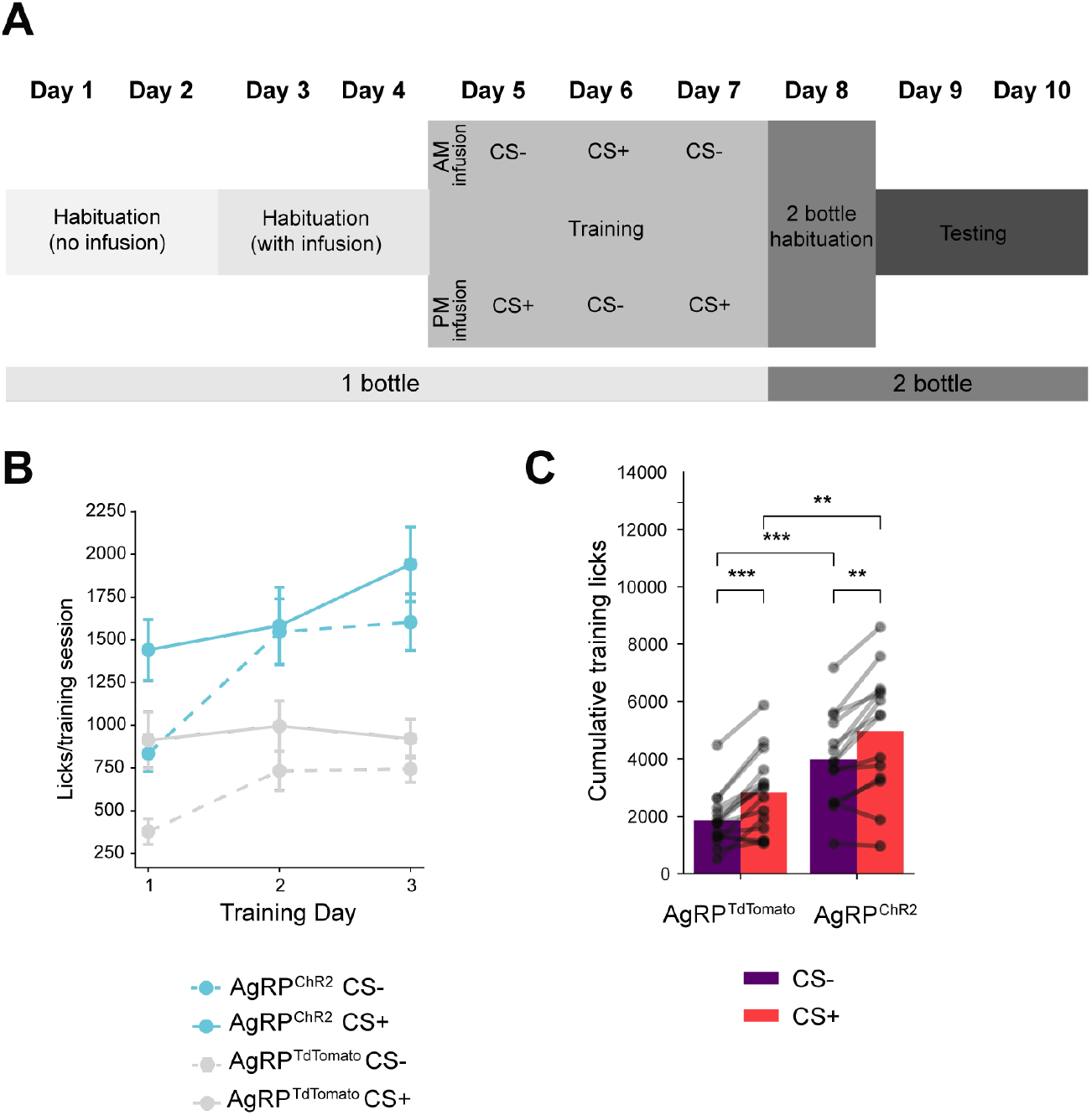
Flavor-nutrient learning protocol and training data. (A) Complete protocol used for FNL. (B) Number of CS+ and CS- licks across training sessions in control (AgRP^tdTomato^) (n=15) and experimental (AgRP^ChR2^) (n=14) mice. ANOVA on Linear Mixed Effects Model (see Supplementary Information for full table). (C) Average number of licks per training session during FNL protocol in control (AgRP^tdTomato^) (n=15) and experimental (AgRP^ChR2^) (n=14) mice. Mixed ANOVA, Chr2: F(1,27)=14.058, p<0.001; CS: F(1,27)=45.376, p<0.0001; Chr2*CS: F(1,27)=0.0007, p=ns. Data are expressed as mean ± SEM. *p<0.05, **p<0.01, ***p<0.001, ****p<0.0001.

**Figure S2.**
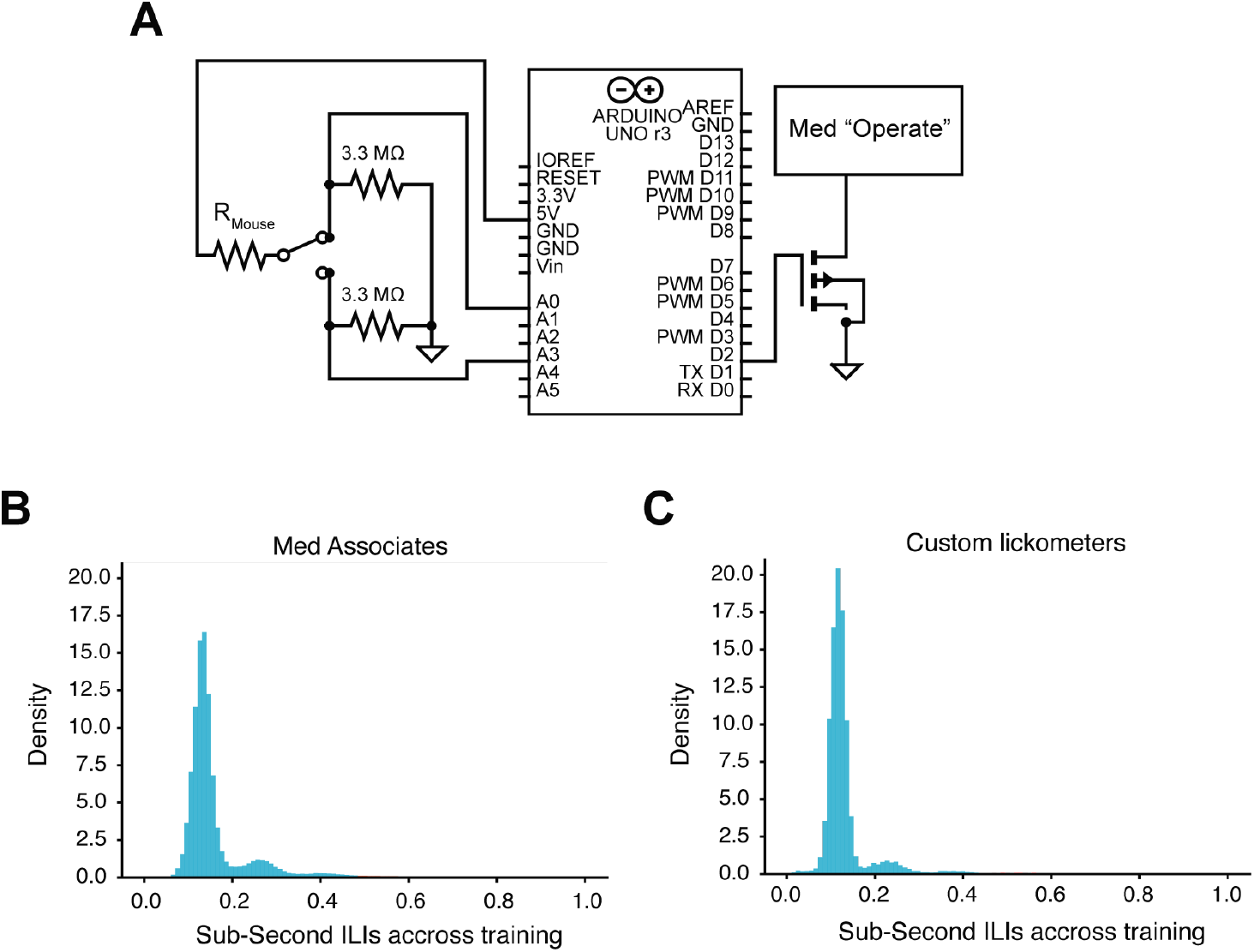
Verification of home-made lickometer equipment. (A) Circuit diagram of custom home-cage lickometer. (B) Inter-lick intervals of n=29 mice measured during FNL training sessions (Figure 1) using commercial Med Associates lickometers. (C) Inter-lick intervals of n=7 mice measured during FNL training sessions during fiber photometry (Figures 4-5) using custom, home-made lickometers.

**Figure S3.**
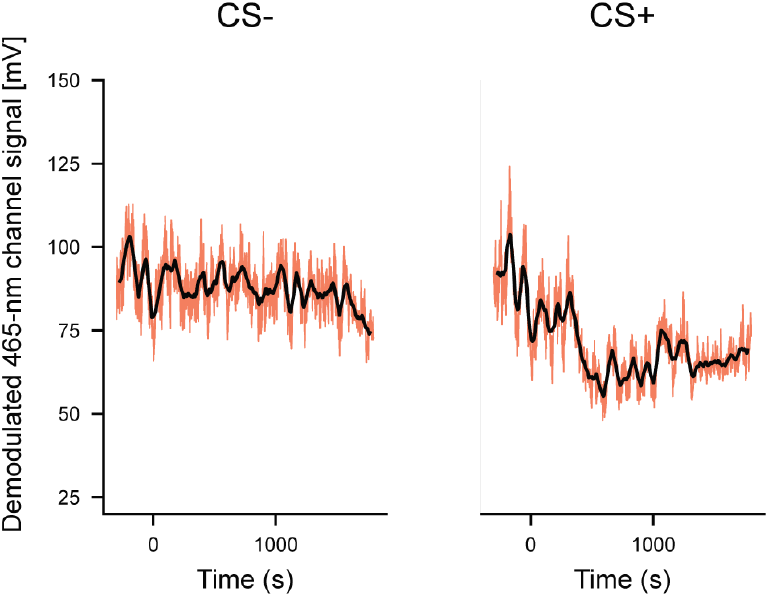
Detrending fiber photometry data for peri-lick bout analyses. Example raw fiber photometry traces with moving averages superimposed. This was used to account for the moving baseline activity when extracting peri-lick bout AgRP neuron activity.

**Figure S4.**
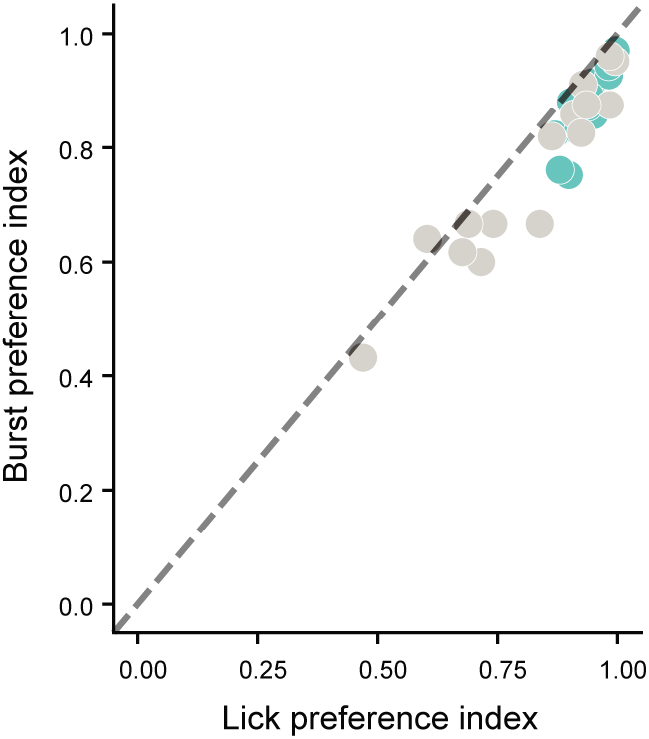
Correlation between lick preference index and burst preference index. The strong correlation between lick preference index and burst preference index justifies the use of the more conservative measure (burst preference index) for modeling (see Methods for additional details). Signed Rank Test, W=7, p<0.0001; Spearman Rank Correlation, ρ=0.907, p<0.0001.

**Figure S5.**
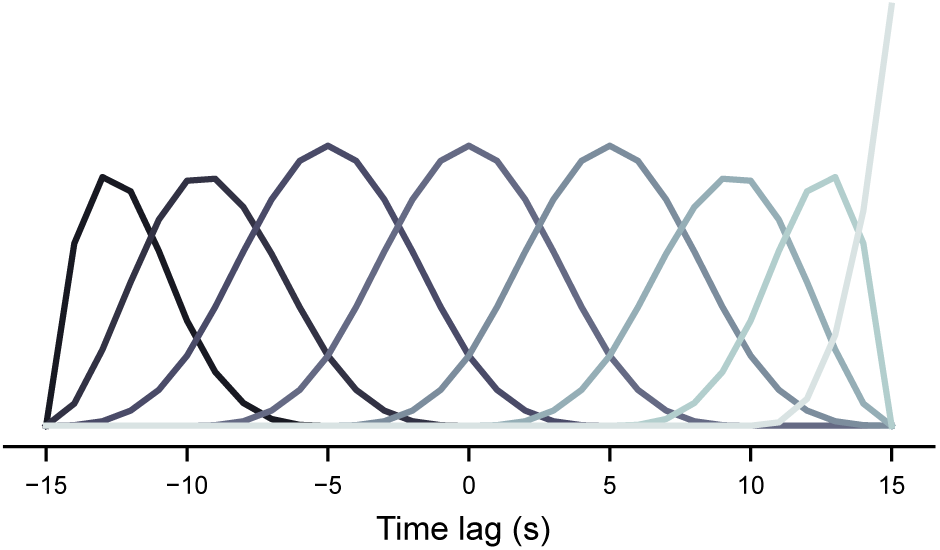
Regression splines for linear mixed effects modeling of peri-lick bout responses. This depicts the basis set of curves used to represent time in mixed effects models of peri-lick bout responses. Model fitting estimates weights for these curves such that their weighted sum reconstructs the mean response for a given CS on a given day. The curves are ordered along the x axis as they are numbered in the regression coefficients. In other words, spline 1 is the left-most curve and spline 8 is the right-most curve.

**Table S1.**
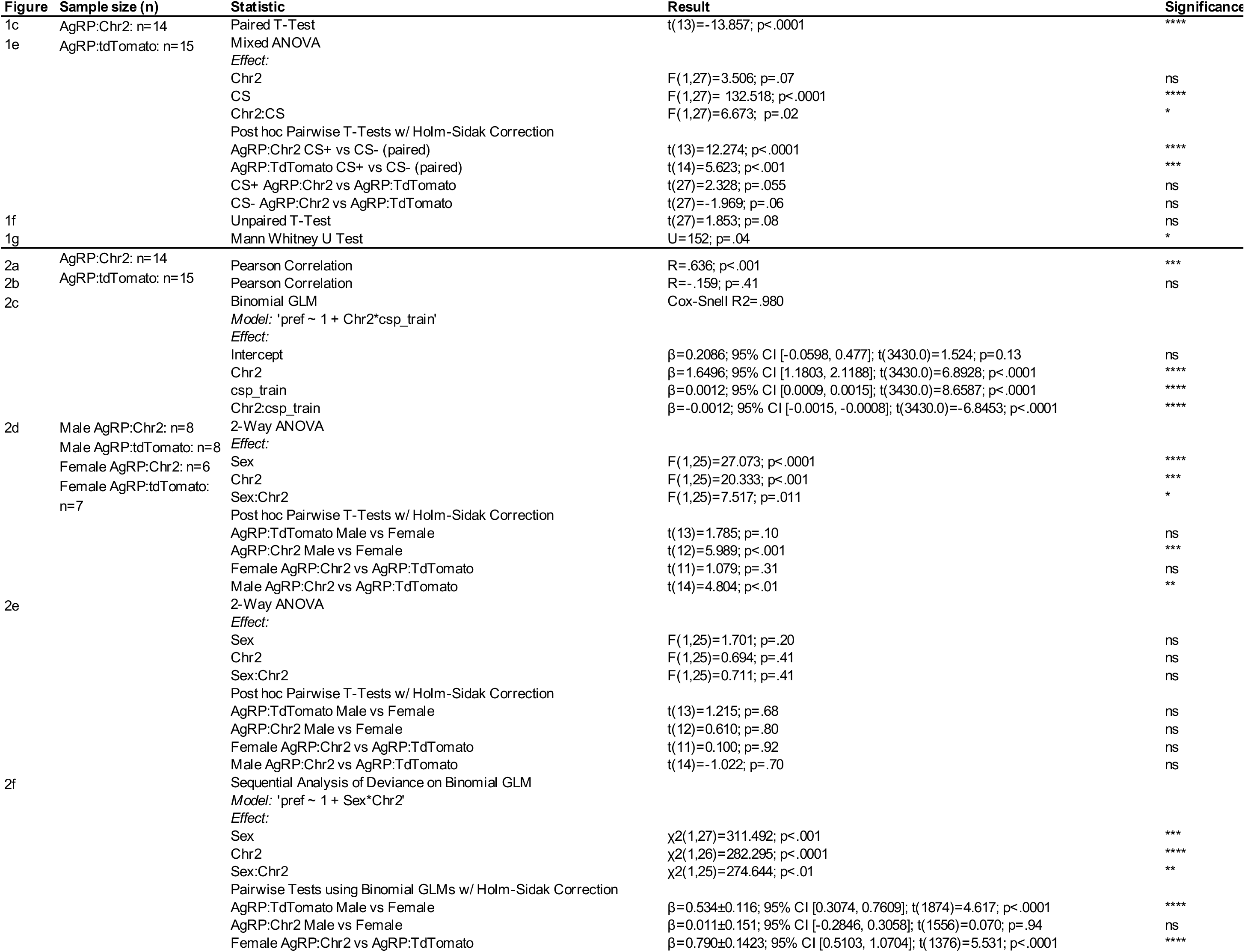
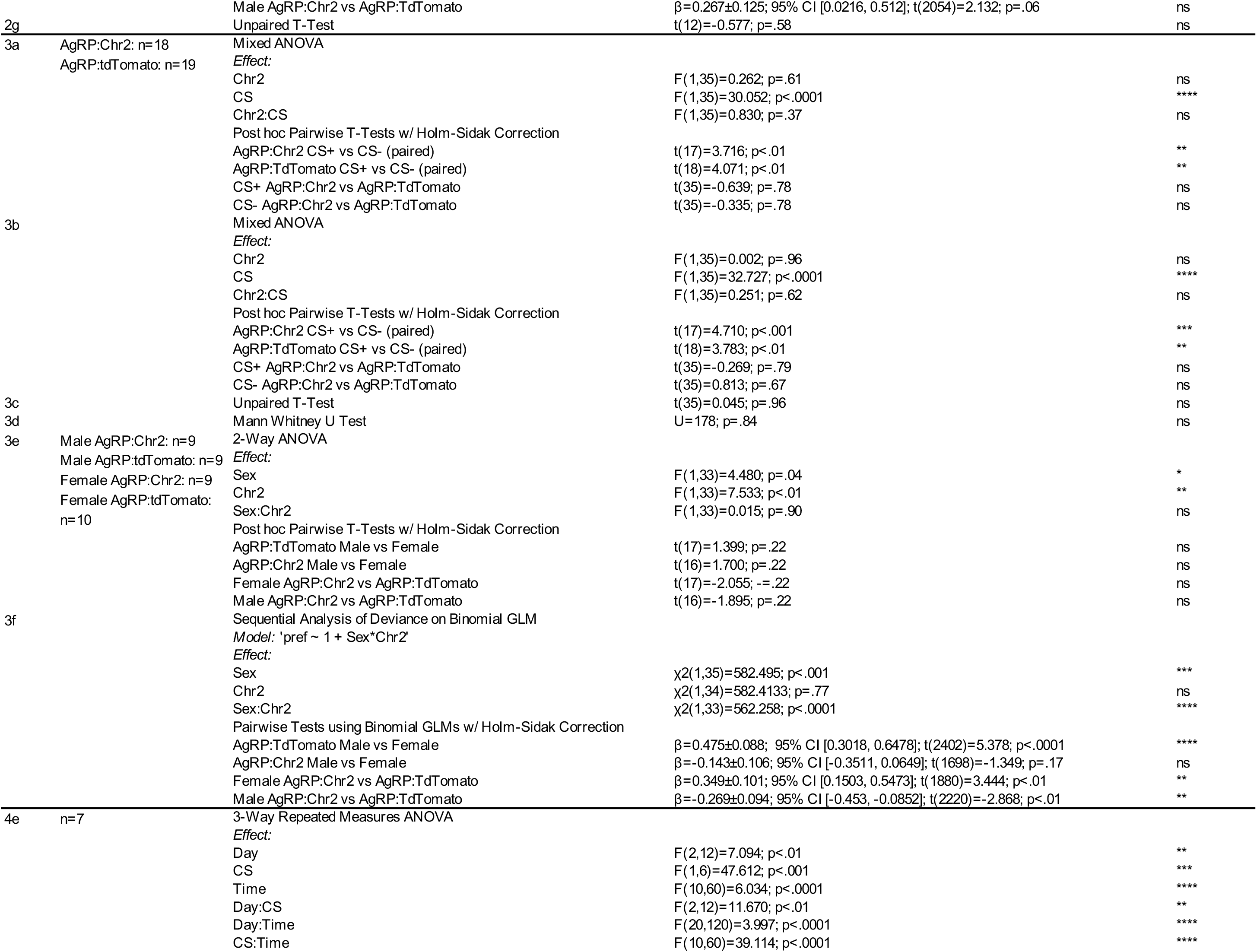
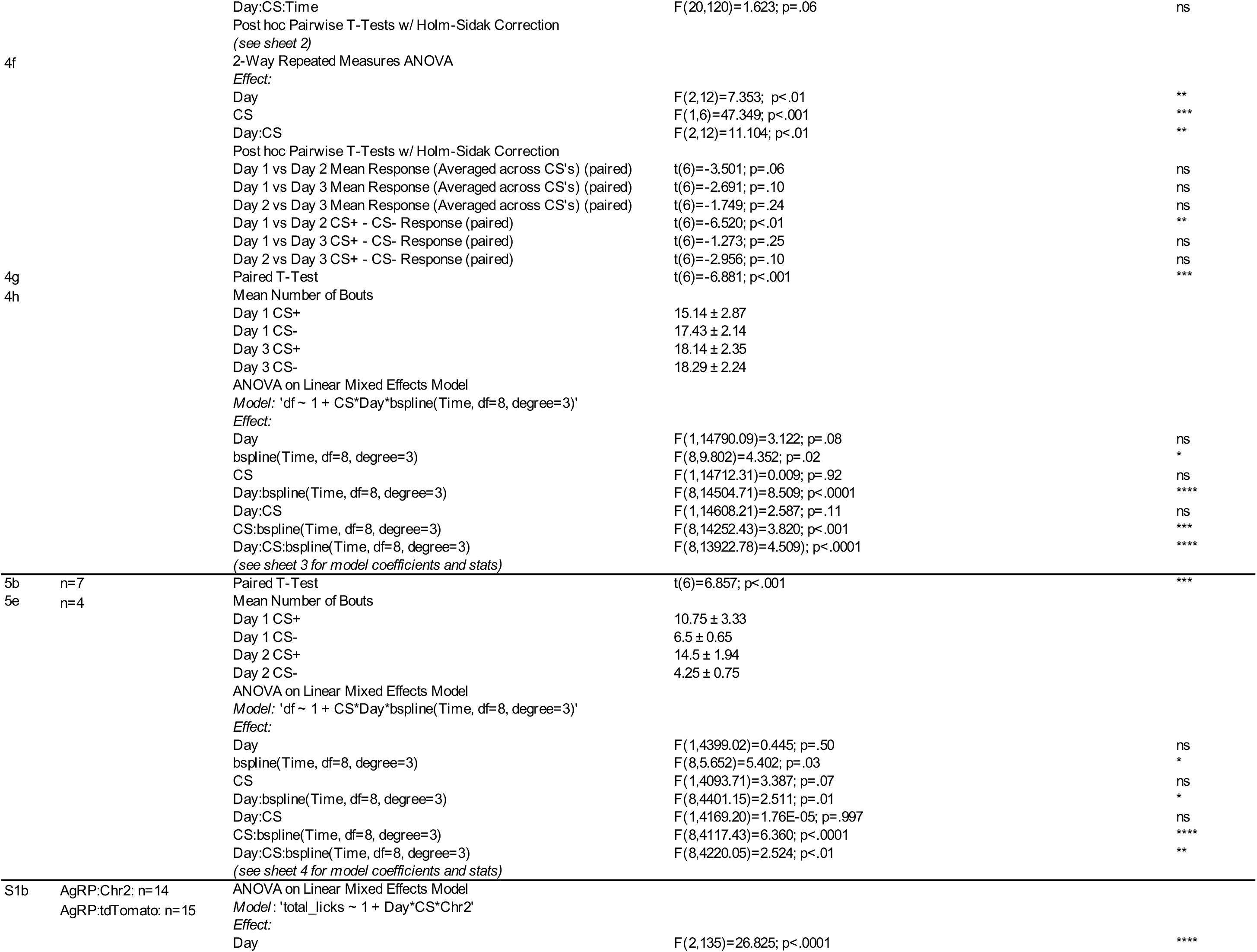
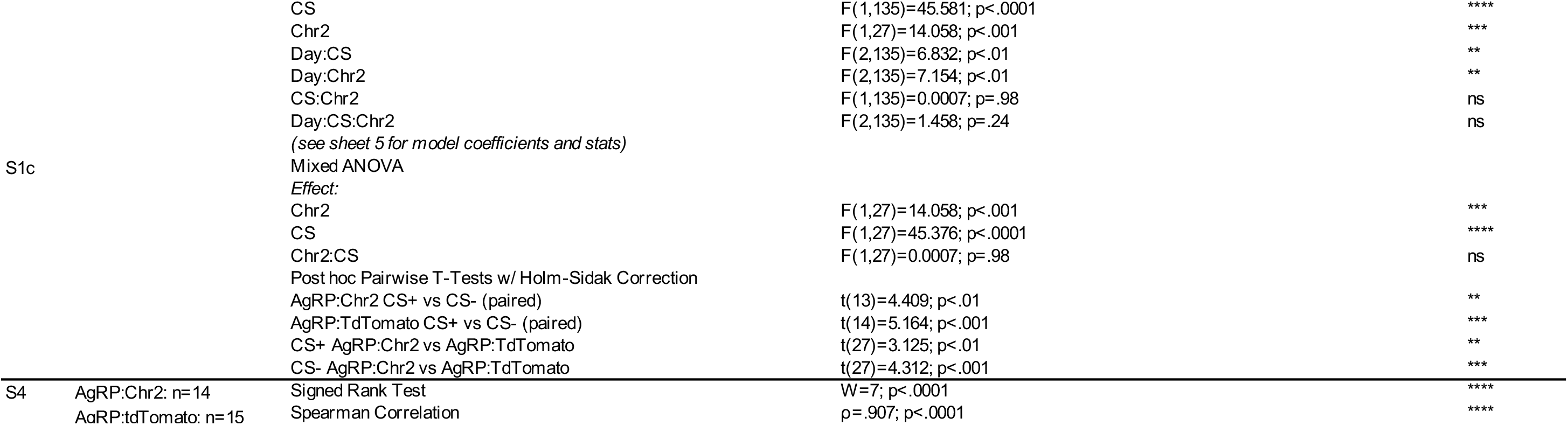

**Table S2.**

| Data (Related to Figure 2) | Sample Size (n) | Statistic | Result | Significance |
| --- | --- | --- | --- | --- |
| Unlimited Intake; Stim Mice Alone | n=14 | Binomial GLM<br>Model: 'pref ~ 1 + csp_train'<br>Effect: | Cox-Snell R2=2.537E-06 |  |
| | | Intercept | $\beta=1.8582$; 95% CI [1.4732, 2.2432]; t(1556.0)=9.4663; p<.0001 | **** |
| | | csp_train | $\beta=0.0$; 95% CI [-0.0002, 0.0002]; t(1556.0)=0.006; p=1.00 | ns |
| Unlimited Intake; Control Mice Alone | n=15 | Binomial GLM<br>Model: 'pref ~ 1 + csp_train'<br>Effect: | Cox-Snell R2=.996 |  |
| | | Intercept | $\beta=0.2086$; 95% CI [-0.0599, 0.4771]; t(1874.0)=1.524; p=0.13 | ns |
| | | csp_train | $\beta=0.0012$; 95% CI [0.0009, 0.0015]; t(1874.0)=8.6587; p<.0001 | **** |
| Unlimited Intake; Male Mice Alone | AgRP:Chr2: n=8<br>AgRP:tdTomato: n=8 | Binomial GLM<br>Model: 'pref ~ 1 + Chr2*csp_train'<br>Effect: | Cox-Snell R2=.992 |  |
| | | Intercept | $\beta=0.0033$; 95% CI [-0.3869, 0.3936]; t(2052.0)=0.0167; p=0.99 | ns |
| | | Chr2 | $\beta=1.3197$; 95% CI [-0.0072, 2.6466]; t(2052.0)=1.9505; p=0.05 | ns |
| | | csp_train | $\beta=0.0014$; 95% CI [0.0011, 0.0018]; t(2052.0)=7.9552; p<.0001 | **** |
| | | Chr2:csp_train | $\beta=-0.0012$; 95% CI [-0.0019, -0.0005]; t(2052.0)=-3.4857; p<.001 | *** |
| Unlimited Intake; Female Mice Alone | AgRP:Chr2: n=6<br>AgRP:tdTomato: n=7 | Binomial GLM<br>Model: 'pref ~ 1 + Chr2*csp_train'<br>Effect: | Cox-Snell R2=.992 |  |
| | | Intercept | $\beta=0.0033$; 95% CI [-0.3869, 0.3936]; t(2052.0)=0.0167; p=0.99 | ns |
| | | Chr2 | $\beta=1.3197$; 95% CI [-0.0072, 2.6466]; t(2052.0)=1.9505; p=0.05 | ns |
| | | csp_train | $\beta=0.0014$; 95% CI [0.0011, 0.0018]; t(2052.0)=7.9552; p<.0001 | **** |
| | | Chr2:csp_train | $\beta=-0.0012$; 95% CI [-0.0019, -0.0005]; t(2052.0)=-3.4857; p<.001 | *** |

